# STAREG: an empirical Bayesian approach to detect replicable spatially variable genes in spatial transcriptomic studies

**DOI:** 10.1101/2023.05.30.542607

**Authors:** Yan Li, Xiang Zhou, Rui Chen, Xianyang Zhang, Hongyuan Cao

## Abstract

Identifying replicable genes that display spatial expression patterns from different yet related spatially resolved transcriptomic studies provides stronger scientific evidence and more powerful inference. We present an empirical Bayesian method, STAREG, for identifying replicable spatially variable genes in data generated from various spatially resolved transcriptomic techniques. STAREG models the joint distribution of *p*-values from different studies with a mixture model and accounts for the heterogeneity of different studies. It provides effective control of the false discovery rate and has higher power by borrowing information across genes and different studies. Moreover, it provides different rankings of important spatially variable genes. With the EM algorithm in combination with pool-adjacent-violator-algorithm (PAVA), STAREG is scalable to datasets with tens of thousands of genes measured on tens of thousands of spatial spots without any tuning parameters. Analyzing three pairs of spatially resolved transcriptomic datasets using STAREG, we show that it makes biological discoveries that otherwise cannot be obtained by using existing methods.

## 1 Introduction

Technological advances in spatially resolved transcriptomics (SRT) have enabled high throughput gene expression profiling while preserving spatial location information of cells in tissues or cell cultures (Asp *et al*., 2020; Ståhl *et al*., 2016; Rodriques *et al*., 2019; Stickels *et al*., 2021). This additional dimension of spatial information brings new perspectives on the cellular transcriptome, allowing researchers to uncover complex cellular and sub-cellular architecture in heterogeneous tissues, which provides crucial insights into complex biological processes (Asp *et al*., 2020). In SRT studies, genes that show spatial expression variations across spatial locations on a tissue section are known as spatially variable genes (SVGs). Detection of SVGs is an important first step in characterizing the spatial transcriptomic landscapes of complex tissues (Hu et al., 2021a).

Various methods have been developed to identify SVGs from SRT data. For instance, Trendsceek (Edsgärd *et al*., 2018) utilizes a marked point process to identify genes with statistically significant spatial expression trends. SpatialDE (Svensson *et al*., 2018) identifies genes with spatial expression patterns and implements spatial gene-clustering, which relates tissue structure and cell-type composition based on the marker genes. SPARK (Sun *et al*., 2020) identifies SVGs through generalized linear spatial models of count data. SPARKX (Zhu *et al*., 2021) uses a non-parametric approach to rapidly and effectively detect spatially expressed genes in large-scale spatial transcriptomic studies. SpaGCN (Hu et al., 2021b) integrates gene expression, spatial location, and histology in SRT data analysis to identify spatial domains with coherent expression and histology and detect genes enriched in the domains. Despite these advances in SVG detection from SRT data, the important issue of replicability of analytical results of SVG detection based on SRT data under distinct cellular environments measured with different techniques has not been addressed.

Replicability is a cornerstone of modern scientific research. Consistent results obtained from different studies with different data provide stronger scientific evidence. In addition, pooling inferences made under different yet related conditions enables researchers to gain statistical power. Statistical definition of replicability can be found in Benjamini et al. (2009); Bogomolov and Heller (2013); Hung and Fithian (2020). In the SVG detection of SRT data, tens of thousands of genes are tested simultaneously. An acute problem is multiple comparison adjustment. A classic approach for multiple comparisons is the false discovery rate (FDR) control procedure proposed in Benjamini and Hochberg (1995), which is known as the BH procedure. When there is more than one study, an *ad hoc* approach to have replicability is to first compute *p*-values from each study, then apply a multiple comparison procedure separately for each study, and finally declare replicable SVGs as the intersection of significant genes from different studies. Not only does this procedure lose power, but it also does not have FDR control, as demonstrated in our simulation studies. Intuitively speaking, if there is no danger that a multiple testing procedure produces false positives, then this *ad hoc* approach would work. However, multiple testing procedures have a non-zero probability of producing false positives unless the procedure does not reject. Thus, an approach that provides control over false positives in each study separately does not guarantee control over false positives to test replicability (Bogomolov and Heller, 2013). As a conservative alternative, researchers may use the maximum of *p*-values from different studies for each gene and implement a multiple testing procedure to claim replicability (Benjamini et al., 2009). This can lose substantial power, as demonstrated in our simulation studies and data analysis.

In this paper, we propose a robust, powerful empirical Bayesian approach for <u>s</u>patial <u>t</u>ranscriptomic <u>a</u>nalysis of <u>r</u>eplicable <u>e</u>xpressed <u>g</u>enes (STAREG) from two SRT studies. Based on the hidden state of whether a gene is an SVG or not, we extend the two-group model to a four-group model (Efron *et al*., 2001; Efron, 2012; Chung et al., 2014). A gene is a replicable SVG if it is significant in both studies. We do not need the two studies to have the same alignment as rotations or normalizations may distort important biological information. In addition, our framework can effectively handle a composite null hypothesis without power loss. A frequentist approach guards against the worst case, while the empirical Bayes approach relies on the prior probability for different hypotheses states (Bogomolov and Heller, 2022). We use local false discovery rate (Lfdr), which is the posterior probability of being null given data, as the test statistic (Efron *et al*., 2001). Lfdr combines the information in both null and non-null hypotheses and thus provides different rankings of importance compared to *p*-value-based methods, such as the BH procedure, which is obtained only under the null hypothesis. A step-up procedure is used to get asymptotic FDR control (Sun and Cai, 2007). By borrowing information across genes and across different studies, STAREG is more powerful at detecting replicable SVGs while controlling the FDR. By combining EM algorithm and pool-adjacent-violator-algorithm (PAVA), STAREG is scalable to datasets with tens of thousands of genes measured on tens of thousands of spatial spots without any tuning parameters (Dempster *et al*., 1977; Robertson et al., 1988; De Leeuw et al., 2009). We apply STAREG to three pairs of SRT datasets collected from distinct technologies to detect replicable SVGs: mouse olfactory bulb data measured with spatial transcriptomics (ST) technology (Ståhl *et al*., 2016), human breast cancer data measured with ST technology (Ståhl *et al*., 2016) and 10X Visium technology, and mouse cerebellum data measured with Slide-seq (Rodriques *et al*., 2019) technology and Slide-seqV2 (Stickels *et al*., 2021) technology. We show that STAREG uncovers important replicable biological discoveries that otherwise cannot be made by existing methods.

## 2 Results

### 2.1 Method overview of STAREG

We describe the workflow of STAREG for replicability analysis of SVG detection from SRT studies. A schematic of STAREG is shown in Fig. 1a. Suppose there are *m* common genes in two SRT datasets. For each SRT dataset, we test *m* hypotheses simultaneously, where the null hypothesis for *i*th gene states that it is not an SVG, and the non-null hypothesis states that it is an SVG. We require *p*-values to have standard uniform distribution under the null for both studies. Any statistical methods that produce well-calibrated *p*-values can be used in this step. STAREG uses paired *p*-values (*p*_1_*_i_, p*_2_*_i_*)*, i* = 1*, . . ., m* as input to the replicability analysis. Let *θ_ji_* denote the hidden state of *i*th hypothesis in study *j* (*j* = 1, 2), where *θ_ji_* = 1 indicates *i*th gene is an SVG in study *j*, and *θ_ji_* = 0 otherwise. Write the joint hidden states across two studies as *τ_i_* ∈ {(0, 0), (0, 1), (1, 0), (1, 1)} with prior probabilities *P* (*τ_i_* = (*k, l*)) = *ξ_kl_*, where *k, l* = 0, 1 and Σ*_k,l_ ξ_kl_* = 1. Our *i*th null hypothesis for replicability of SVG in two studies is *H_i_*_0_ : *τ_i_* ∈ {(0, 0), (0, 1), (1, 0)}, *i* = 1*, . . ., m.* We use a four-group model for the joint distribution of (*p_i_*_1_*, p_i_*_2_)*, i* = 1*, . . ., m,* where the distribution of *p*-values under the null is assumed to be standard uniform for both studies, and *p*-value distributions under the non-null are estimated non-parametrically and can be different in two SRT studies to accommodate heterogeneity. For *i*th gene, we use Lfdr as our test statistic, which is defined as the posterior probability of *H_i_*_0_ given (*p_i_*_1_*, p_i_*_2_)*, i* = 1*, . . ., m.* We use the EM algorithm in combination with PAVA to estimate unknown parameters (*ξ*_00_*, ξ*_01_*, ξ*_10_*, ξ*_11_) and *p*-value distributions under the non-null for study 1 and study 2 non-parametrically. Denote estimated Lfdr as 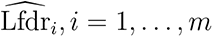. Small Lfdr indicates strong evidence against the null. The rejection region can be written as 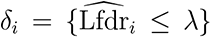. We implement a step-up procedure based on 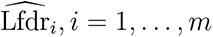 to identify replicable SVGs from the two SRT studies (Sun and Cai, 2007). More details of STAREG can be found in the Methods section and Supplementary Materials.

**Figure 1:**
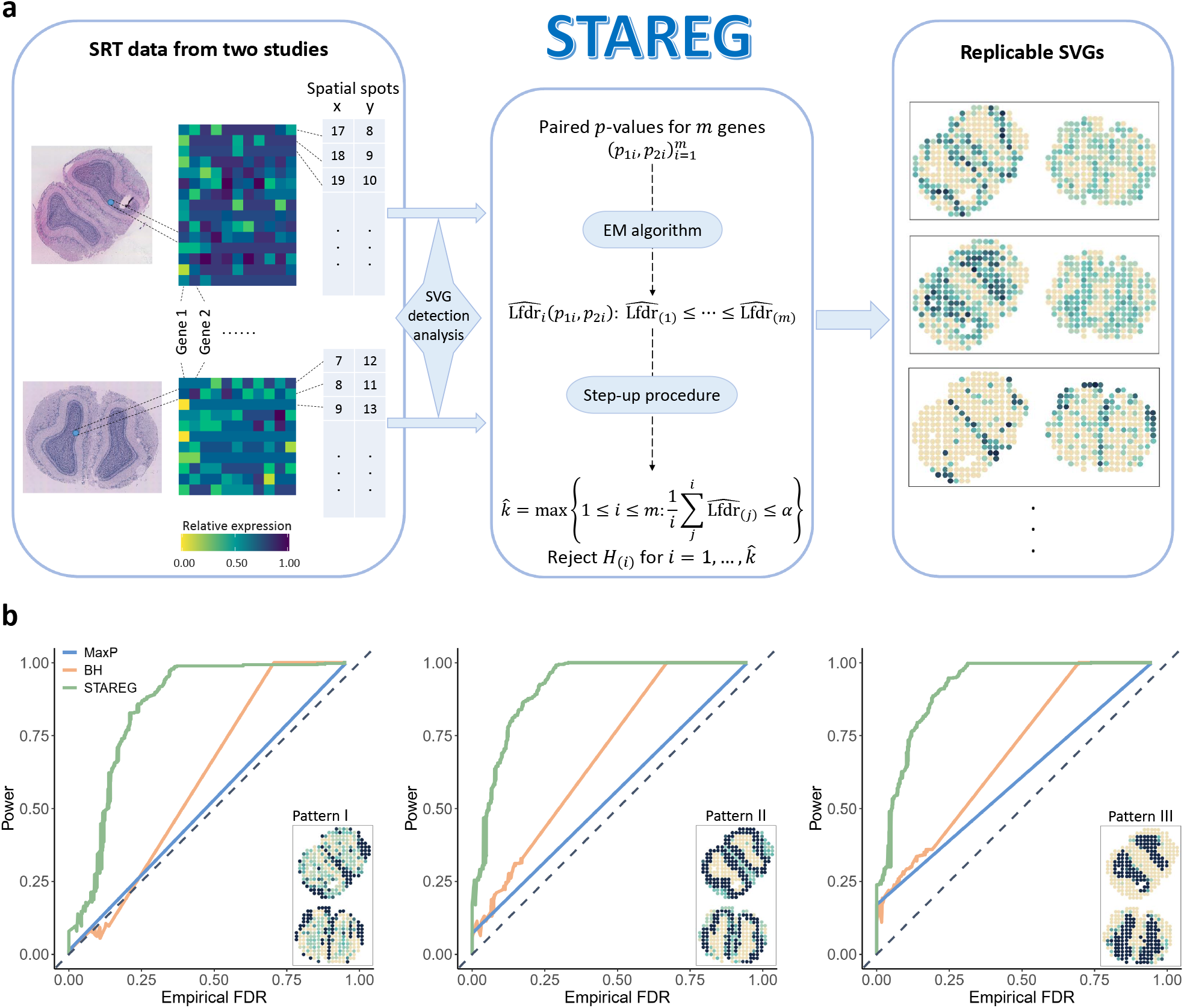
a. Schematic of the STAREG. **b** Plots of power (y-axis) and empirical FDR (x-axis) of different methods from realistic simulation studies based on the mouse olfactory bulb data. Three spatial expression patterns used for simulation studies are labeled inside corresponding panels.

### 2.2 Simulation studies

We conducted realistic simulation studies to evaluate FDR control and the power of STAREG. FDR is defined as the expectation of the number of false rejections over the total number of rejections. Power refers to the true positive rate, which is the expectation of the number of replicable findings over the total number of non-null hypotheses. Specifically, data were generated based on two replicates of the mouse olfactory bulb data measured with ST technology (Ståhl *et al*., 2016) with parameters inferred from SPARK (Sun *et al*., 2020). Details of data generation were provided in Supplementary Materials. In addition to STAREG, we considered two other replicability procedures, BH and MaxP, for comparison. Detailed implementations of these two methods can be found in the Methods section.

Simulated SRT data corresponding to three spatial expression patterns as in Fig. 1b were generated in pairs and analyzed separately with SPARK to produce input for replicability analysis. We observe that the FDR of STAREG and MaxP can be controlled at a nominal level, while BH failed to control FDR for the replicability null hypothesis in some settings (Supplementary Fig. S1a, S2). At the same FDR level, STAREG demonstrated substantial power gain across various settings (Supplementary Fig. S1a, S3). To have a fair comparison, we calculated empirical FDR and power with a range of FDR cutoffs based on different methods in Fig. 1b and Supplementary Fig. S1b, S4. The number of genes was set as *m* = 10, 000, and prior probabilities used to produce four hidden states were set as *ξ*_00_ = 0.9*, ξ*_01_ = *ξ*_10_ = 0.025, and *ξ*_11_ = 0.05. Overall, the simulation results suggest that STAREG has superior performance to competing methods. The reason for the favorable performance of STAREG is due to the structure of the test statistic Lfdr, which efficiently combines information in both null and non-null and provides a better ranking of important genes compared with *p*-value-based methods, where only information contained in the null hypothesis is utilized. The heterogeneity of different studies manifest through *ξ*_01_ and *ξ*_10_. We model such heterogeneity through a four-group model expanding classic two-group model (Efron *et al*., 2001).

We evaluated the computational time of STAREG in simulation studies. As any statistical methods that produce well-calibrated *p*-values can be used for each SRT study, and all methods we compare work directly with paired *p*-values, we only considered the computational overhead of replicability analysis after obtaining paired *p*-values from simulated SRT data. Prior probabilities used to produce four hidden states were set as *ξ*_00_ = 0.9*, ξ*_01_ = *ξ*_10_ = 0.025, and *ξ*_11_ = 0.05. Table 1 summarizes the computational time of different methods for SRT data with different numbers of genes. Computations were carried out in an Intel(R) Core(TM) i7-9750H 2.6GHz CPU with 64.0 GB RAM laptop. We observe that all methods are quick to compute. STAREG takes longer time than BH and MaxP, though such differences can be ignored in practical data analysis. More details on simulation studies can be found in Supplementary Materials.

**Table 1:** Computational time (in seconds) for replicability analysis in simulation studies

| # of genes | Pattern I |  |  | Pattern II |  |  | Pattern III |  |  |
| --- | --- | --- | --- | --- | --- | --- | --- | --- | --- |
|  | BH | MaxP | STAREG | BH | MaxP | STAREG | BH | MaxP | STAREG |
| 5,000 | 0.005 | 0.012 | 1.246 | 0.005 | 0.011 | 0.990 | 0.005 | 0.014 | 0.931 |
| 10,000 | 0.007 | 0.020 | 4.872 | 0.007 | 0.018 | 3.806 | 0.006 | 0.026 | 4.053 |
| 20,000 | 0.013 | 0.038 | 19.147 | 0.008 | 0.033 | 14.072 | 0.007 | 0.034 | 13.072 |

### 2.3 Application to mouse olfactory bulb data

We first performed replicability analysis on Replicate 1 and Replicate 8 of mouse olfactory bulb ST data (Ståhl *et al*., 2016) obtained from the Spatial Research Website. Replicate 1 data consist of 10, 373 genes on 265 spots and Replicate 8 data contain 9, 671 genes on 232 spots. We applied SPARK separately on the two datasets to conduct SVG detection analysis, yielding corresponding *p*-values. We next took 9, 329 pairs of *p*-values of common genes as input for replicability analysis.

At FDR level of 0.05, STAREG detected 1, 175 replicable SVGs, including all replicable SVGs identified by BH and MaxP. MaxP identified 559 replicable SVGs, which were also detected by BH (618 replicable SVGs). This is consistent with simulation results that STAREG has higher power. We plot the non-parametric estimates of *p*-value density functions under non-null for two studies in Supplementary Fig. S5a. To assess the quality of the 1, 175 replicable SVGs identified by STAREG, we clustered genes into three groups with distinct spatial expression patterns using R package *amap* v0.8-18. As shown in Fig. 2a and Supplementary Fig. S6a,b, the three distinct spatial patterns in the two studies are consistent and can be matched to three main layers in mouse olfactory bulb: Pattern I corresponds to the glomerular cell layer, Pattern II corresponds to the mitral cell layer and Pattern III corresponds to the granular cell layer. The three patterns can be visualized via three representative genes only detected by STAREG: *Dcn*, *Tspan9* and *Srrm4* (Fig. 2b). We listed spatial expression patterns of 20 randomly selected replicable SVGs only identified by STAREG in two studies as additional evidence of replicability (Supplementary Fig. S6d,e). We also calculated Moran’s *I* statistics (Moran, 1950), a commonly used metric to quantify spatial autocorrelations. The 557 uniquely identified replicable SVGs have larger Moran’s *I* statistic than that of overall 9, 329 genes (Fig. 2f).

**Figure 2:**
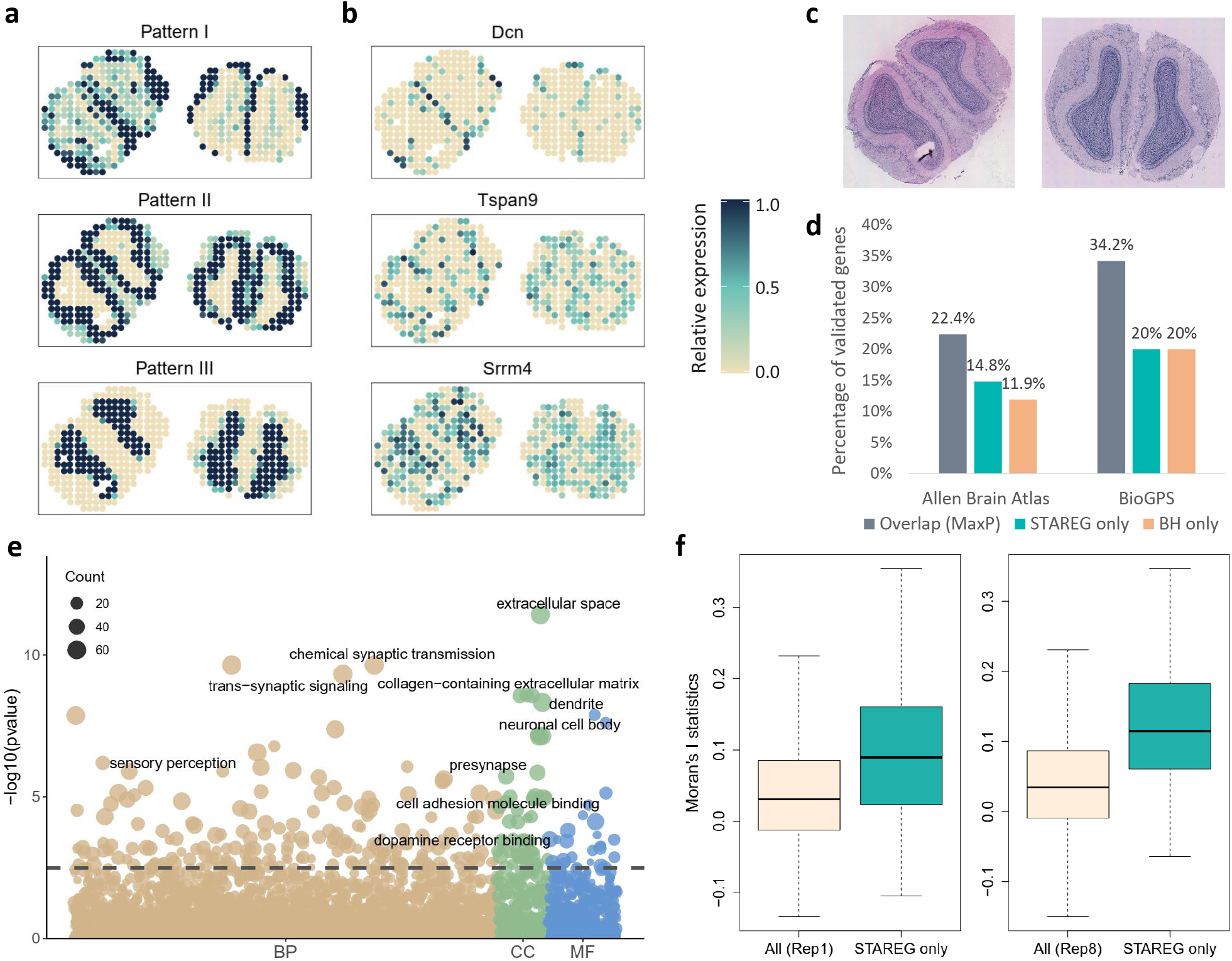
Analysis and validation results of the mouse olfactory bulb data measured with ST technology. **a** Three clustered distinct spatial expression patterns in two SRT studies summarized on the basis of 1, 175 replicable SVGs identified by STAREG (left: Replicate 1; right: Replicate 8). Color represents relative gene expression levels (antique white: low; navy blue: high). **b** Spatial expression patterns of three representative replicable SVGs only identified by STAREG, corresponding to three main cell layers in mouse olfactory bulb (left: Replicate 1; right: Replicate 8). **c** HE images represent the mouse olfactory bulb sections corresponding to Replicate 1 (left) and Replicate 8 (right). **d** Bar chart shows the percentage of replicable SVGs additionally detected by STAREG and BH compared to MaxP that were validated in two reference gene lists from the Harmonizome database: one from the Allen Brain Atlas and the other from the BioGPS. **e** Bubble plot shows *−* log_10_(*p*) of GO terms enriched by the 557 replicable SVGs only identified by STAREG. The dashed line represents an FDR cutoff of 0.05. Color shows different GO term categories: biological process (BP, tan), cellular component (CC, green), and molecular function (MF, blue). The size of bubbles represents counts of corresponding gene sets. **f**Box plot shows Moran’s *I* values of replicable SVGs only identified by STAREG compared to overall genes.

We summarized three lists of reference genes that are related to olfactory bulb from the original ST study (Ståhl *et al*., 2016) and the Harmonizome database (Rouillard *et al*., 2016) to validate replicable SVGs identified by different methods. First, MaxP and BH detected 7 of the 10 genes published in the original ST study (Ståhl *et al*., 2016) that are highly expressed in the mitral cell layer of olfactory bulb, whereas STAREG identified additional 2 such highlighted SVGs. The second gene set from the Allen Brain Atlas (Sunkin *et al*., 2013) contains 1, 894 genes differentially expressed in main olfactory bulb relative to other tissues. Compared to MaxP (22.4% validated), 14.8% of the 616 replicable SVGs only detected by STAREG were in the list, whereas 11.9% of the 59 additional findings by BH were validated. The third gene list consists of 2, 031 genes related to mouse olfactory bulb from BioGPS (Wu *et al*., 2013). Compared to MaxP (34.2% validated), 124 of the 616 SVGs only detected by STAREG were validated, and 12 of the 59 SVGs only identified by BH were in the same list. These validation results again demonstrate the higher power of STAREG.

Finally, we performed Gene Ontology (GO) enrichment analysis and Kyoto Encyclopedia of Genes and Genomes (KEGG) pathway analysis on the 557 replicable SVGs only detected by STAREG to gain additional biological insights (Fig. 2e and Supplementary Fig. S6c). At the FDR level of 0.05, 245 GO terms and 11 KEGG pathways were additionally enriched by STAREG, in which many are related to olfactory bulb development. For example, the GO term of olfactory lobe development with ID GO:0021988, only enriched by STAREG is closely related to the olfactory bulb functions; the GO terms with IDs GO:0035256 and GO:0045744 regulate G protein-coupled receptor signaling pathway, which plays a critical role in olfactory signal transduction, neuronal morphology and axon guidance of mouse (Ebrahimi and Chess, 1998); and the KEGG pathway of oxytocin signaling (mmu04921) is closely related to the modulation of olfactory processing (Adan *et al*., 1995). These additional enrichments in GO terms and KEGG pathways demonstrate the biological significance of findings only detected by STAREG.

### 2.4 Application to human breast cancer data

We performed replicability analysis of human breast cancer data measured with ST (Ståhl *et al*., 2016) and 10X Visium, consisting of 5, 262 genes on 250 spots (ST) and 11, 993 genes on 3, 798 spots (10X Visium), respectively. We analyzed these two datasets with SPARK to produce two sequences of *p*-values. Matched by genes, 4, 943 pairs of *p*-values were obtained as input for replicability analysis.

At the FDR level of 0.05, STAREG identified 566 replicable SVGs, including all 504 SVGs claimed by BH. MaxP detected 500 replicable SVGs, all of which were also identified by BH. We plot the non-parametric estimates of *p*-value density functions under non-null for two studies in Supplementary Fig. S5b.To examine the validity of replicable SVGs identified by STAREG, we clustered the 566 SVGs into two and three groups and summarized the spatial expression patterns based on the ST data and 10X Visium data, respectively (Fig. 3a,b, and Supplementary Fig. S7a,b). For the ST data, Pattern I corresponds to potential tumor regions presented in the HE image (Fig. 3a: top right). For the 10X Visium data, we annotated the corresponding HE image with four cell clusters produced by the t-distributed stochastic neighbor embedding (t-SNE) algorithm (van der Maaten and Hinton, 2008) (Fig. 3b: bottom right). It can be seen that the three patterns are consistent with different cell clusters. The spatial patterns in both studies can be well visualized via a representative gene *RPL27*, which was only identified by STAREG (bottom right of Fig. 3a and bottom left of Fig. 3b). Spatial expression patterns of 20 genes randomly selected from the 62 replicable SVGs only identified by STAREG were also presented (Supplementary Fig. S7d,e), most of which can be validated in corresponding HE images. We further calculated Moran’s *I* statistics (Moran, 1950) to evaluate spatial autocorrelations of the 62 replicable SVGs only identified by STAREG and compared them with baseline Moran’s *I* statistics of all 4, 943 genes. The box plots in Fig. 3e display results based on the ST data (left) and the 10X Visium data (right). It can be seen that the replicable SVGs additionally identified by STAREG have higher Moran’s *I*.

**Figure 3:**
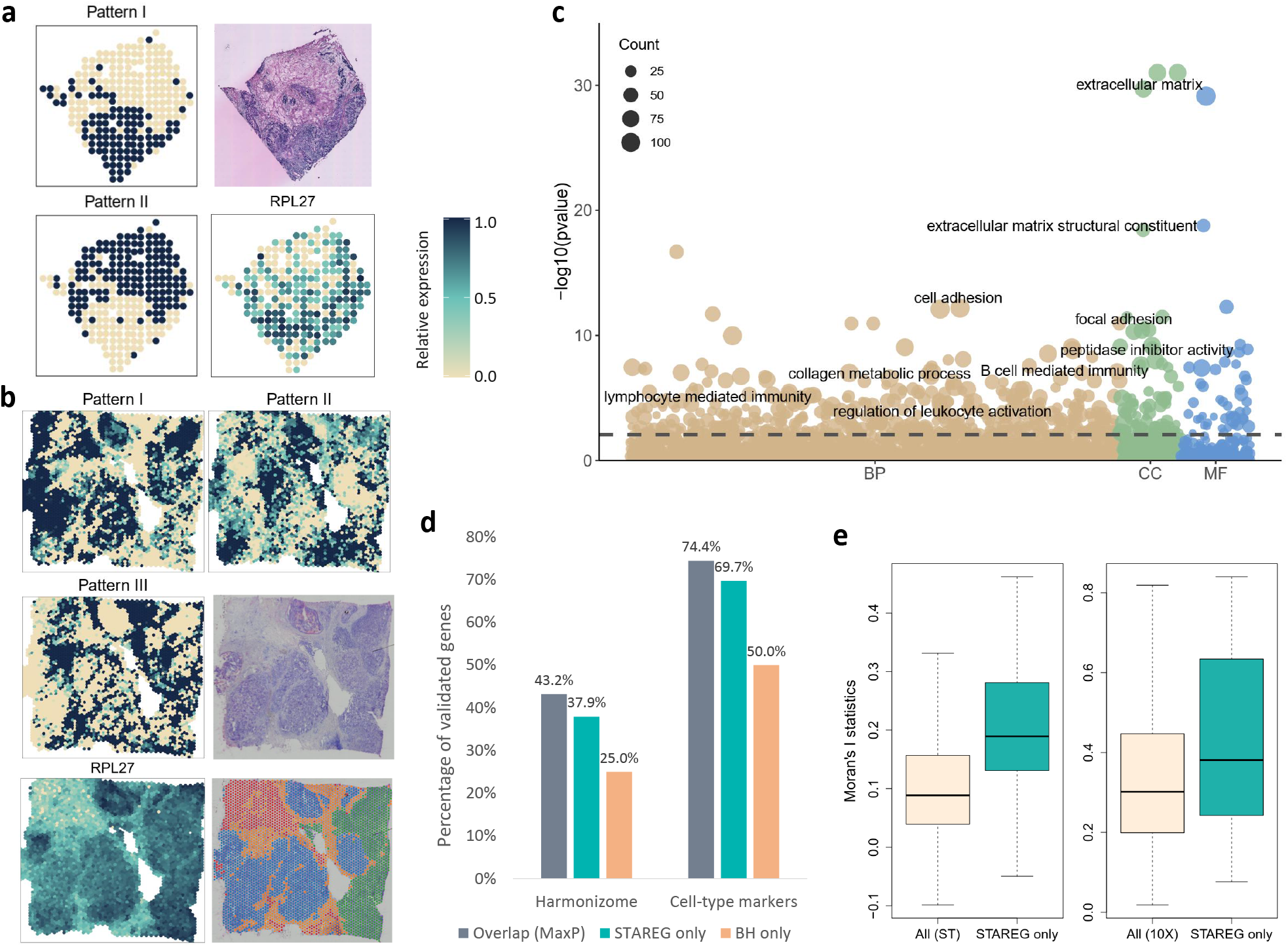
Analysis and validation results of the human breast cancer data measured with ST technology and 10X Visium technology. **a** Pattern I-II summarizes spatial expression patterns in the ST data based on 566 replicable SVGs identified by STAREG. The corresponding HE image (top right) was produced by Ståhl *et al*. (2016), with dark staining representing possible tumor regions. The spatial expression pattern of a representative gene only identified by STAREG (*RPL27*) is presented at the top right. Color represents relative gene expression levels (antique white: low; navy blue: high). **b** Pattern I-III show three clusters summarized in the 10X Visium data based on 566 replicable SVGs identified by STAREG. Corresponding HE image and annotations of cell clusters produced by t-SNE obtained from the 10X Visium repository are presented at the bottom right. The spatial expression pattern of gene *RPL27* is shown at the bottom left. **c** Bubble plot shows *−* log_10_(*p*) of GO terms enriched by 566 replicable SVGs identified by STAREG. The dashed line represents an FDR cutoff of 0.05. The color shows different GO term categories: BP (tan), CC (green), and MF (blue). Bubble size represents the count of the corresponding gene set. **d** Bar chart shows the percentage of replicable SVGs identified by different methods that were validated in two reference gene lists related to human breast cancer: one from the Harmonizome database; the other from Wu *et al*. (2021). **e** Box plot of Moran’s *I* statistics for replicable SVGs only detected by STAREG compared to all genes based on the ST study and the 10X Visium study.

We used two lists of publicly available breast cancer-related genes to validate our results. First, we validated our results in a list of 3, 505 genes that are relevant to breast cancer obtained from the Harmonizome database (Rouillard *et al*., 2016). As in the left panel of Fig. 3d, 43.2% of the 500 SVGs identified by MaxP are in the gene list. Compared to MaxP, 37.9% of the replicable SVGs uniquely identified by STAREG and 1 of the 4 SVGs additionally identified by BH were validated in the same list. Second, we obtained a list of 4, 842 celltype marker genes in human breast cancer (Wu *et al*., 2021) to validate our results. Again, compared to MaxP (74.4% validated), 69.7% of replicable SVGs only detected by STAREG were validated, whereas only 2 of the 4 SVGs uniquely identified by BH are in the same list. Overall, the validation results provide additional evidence of biological findings by STAREG in the replicability analysis.

We finally conducted functional enrichment analysis on the replicable SVGs discovered by BH and STAREG (Fig. 3c and Supplementary Fig. S7c). At FDR level 0.05, BH enriched 532 GO terms and 21 KEGG pathways, whereas STAREG enriched 554 GO terms and 25 KEGG pathways. In addition to overlaps with BH, 65 GO enrichment terms and 4 KEGG pathways were only generated by STAREG, many of which are closely related to breast cancer development. For example, several GO terms related to immune system development (GO:0002520, GO:0140375) were only enriched by STAREG, and the vital role of the immune system played in breast cancer initiation and development has been recognized (Emens, 2012; Gonzalez et al., 2018); the GO terms GO:0032635 and GO:0032675 enriched only by STAREG regulate the production of interleukin-6, whose signaling pathway plays a significant role in immunopathogenesis and treatment of breast cancer (Masjedi *et al*., 2018); and the KEGG pathway of focal adhesion (hsa04510) is a key factor of breast cancer risk (Sahana *et al*., 2021).

### 2.5 Application to mouse cerebellum data

We analyzed two large-scale SRT datasets measured in mouse cerebellum: a Slide-seq dataset containing 18, 082 genes on 28, 352 beads and a Slide-seqV2 dataset consisting of 20, 117 genes on 11, 626 beads. Due to the computational complexity of these datasets, we applied SPARKX (Zhu *et al*., 2021) to analyze them separately, resulting in two sequences of *p*-values from corresponding studies. By intersecting genes in these two studies, we performed replicability analysis of SVG detection on 16, 873 pairs of *p*-values.

At the FDR cutoff of 0.05, MaxP identified 286 replicable SVGs, all of which were identified by BH and STAREG. The 844 replicable SVGs identified by STAREG include all 476 replicable SVGs identified by BH. We plot the non-parametric estimates of *p*-value density functions under non-null for two studies in Supplementary Fig. S5c. Based on these two datasets, we clustered the 844 replicable SVGs identified by STAREG into three groups and summarized distinct spatial expression patterns (Fig. 4a). We observe that consistent spatial patterns were shown in the two studies, with Pattern I corresponding to the spatial distribution of the granular cell layer, Pattern III corresponding to the purkinje cell layer, and Pattern I corresponding to other cell layers. STAREG uniquely identified several well-known cell type marker genes in mouse cerebellum, such as *Ppp1r17* (Endo et al., 1999), *Gabra1* (Kato, 1990), *Edil3* (Ito, 2012), *Gdf10* (Mecklenburg et al., 2014), *Ptprk* and *Nxph1* (Allen brain atlas). We listed the spatial expression pattern for two replicable SVGs only identified by STAREG, *Meg3* and *Ppp1r17*, based on the Slide-seq data and Slide-seqV2 data as examples of the granular layer and purkinje layer in mouse cerebellum (Fig. 4b). More examples can be found in Supplementary Fig. S8b,c. Spatial autocorrelations of replicable genes additionally detected by STAREG were further demonstrated by Moran’s *I* statistics (Moran, 1950) (Fig. 4d).

**Figure 4:**
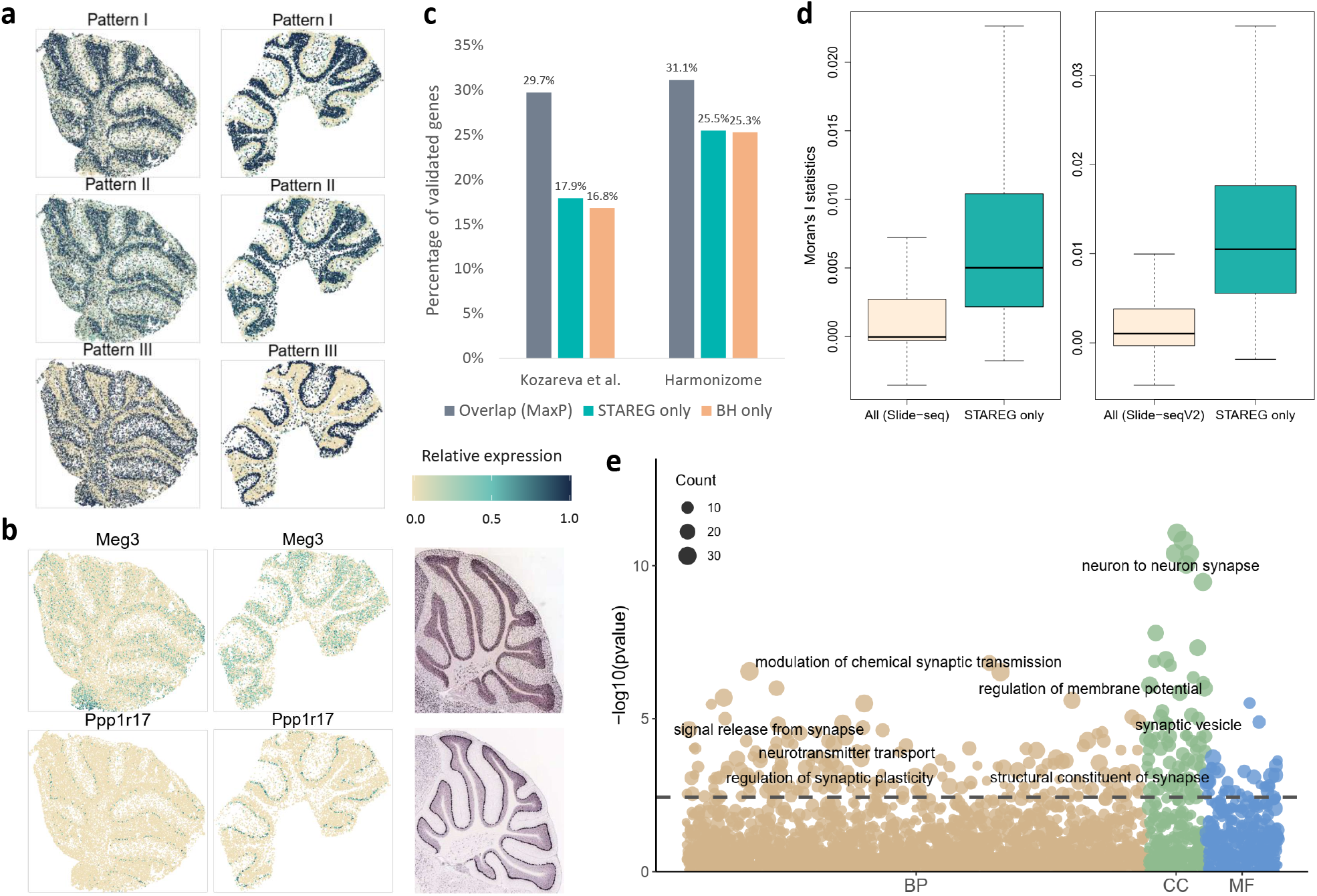
Analysis and validation results of the mouse cerebellum data measured with Slideseq technology and Slide-seqV2 technology. **a** Three spatial expression patterns summarized in the Slide-seq dataset (left) and the Slide-seqV2 dataset (right) based on 844 replicable SVGs identified by STAREG. Color represents relative expression levels (antique white: low; navy blue: high). **b** Spatial expression patterns of two genes only identified by STAREG based on the Slide-seq data (left) and the Slide-seqV2 data (middle). The in situ hybridization images of corresponding genes obtained from the Allen Brain Atlas are presented right. **c** Bar chart displays the percentage of replicable SVGs identified by different methods validated in two published reference gene lists related to mouse cerebellum: one from Kozareva et al. (2021) and the other from the Harmonizome database. **d** Box plot of Moran’s *I* statistics for the 368 replicable SVGs only detected by STAREG based on the Slide-seq data (left) and the Slide-seqV2 data (right). **e** Bubble plot shows *−* log_10_(*p*) of GO terms enriched by the 368 replicable SVGs only identified by STAREG. The dashed line represents the FDR cutoff of 0.05. The color shows different GO term categories: BP (tan), CC (green), and MF (blue). The size of each bubble represents the count of the corresponding gene set.

We validated the quality of replicable SVGs identified by STAREG using two lists of genes related to mouse cerebellum that were published in previous literature. First, a list of 995 highly differentially expressed genes across all cell clusters in mouse cerebellar cortex (Kozareva et al., 2021) were obtained to validate the results. 17.9% of the 558 replicable SVGs identified by STAREG but not detected by MaxP are in the reference gene list, whereas 16.8% of the 190 replicable SVGs detected by BH but not by MaxP were validated in the same list (Fig 4c: left). Second, we obtained a list of 2, 431 genes that were differentially expressed in the cerebellum from the Harmonizome database. Compared to MaxP (31.1% validated), 25.5% of the replicable SVGs only detected by STAREG were successfully validated, whereas 25.3% of the replicable SVGs additionally identified by BH were in the same list. These validation results provide additional biological evidence for the favorable performance of STAREG. Additionally, we performed GO and KEGG enrichment analysis on the 368 replicable SVGs only identified by STAREG to examine the additional biological findings. At the FDR level of 0.05, the 368 uniquely identified genes enriched 271 GO terms and 24 KEGG pathways (Fig. 4e and Supplementary Fig. S8a). Many of them are directly related to the generation and organization of cerebellum structure, such as GO terms of the neuron-to-neuron synapse (GO:0098984), synapse assembly (GO:1904861, GO:0007416), regulation of synapse organization, structure or activity (GO:0050807, GO:0050803), parallel fiber to Purkinje cell synapse (GO:0098686), and pathways of neurodegeneration - multiple diseases (mmu05022). These additional biologically relevant findings make STAREG desirable for replicability analysis of SVG detection in SRT studies.

## 3 Discussion

In this paper, we present STAREG, an empirical Bayesian approach for the detection of replicable SVGs in SRT data across two studies. By borrowing information across genes and different studies, STAREG has higher statistical power while maintaining asymptotic FDR control. We conducted extensive simulation studies to demonstrate FDR control and power gain of STAREG over two competing methods (MaxP and BH). Consistent with simulation studies, analysis results of SRT data from different species, regions, and tissues generated by different technologies demonstrate the favorable performance of STAREG. Important biological findings are revealed by STAREG, which otherwise cannot be obtained by using existing methods.

We mainly applied SPARK (Sun *et al*., 2020) or SPARK-X (Zhu *et al*., 2021) to obtain *p*-values from individual SRT studies. STAREG is versatile, platform-independent, and only needs *p*-values as input. We require *p*-values to be uniformly distributed under the null. Such a requirement is needed in other methods, such as BH and MaxP. In principle, any method that produces well-calibrated *p*-values can be used. In addition, we do not require two tissue sections in SRT studies to have the same spatial expression patterns, making STAREG applicable for SRT data with different sizes, resolutions, and alignments obtained from different technologies. In practice, using alignments, rotations and normalizations may distort important biological information contained in the data.

STAREG uses Lfdr based on a four-group model as the test statistic to efficiently accommodate the composite null hypothesis in replicability analysis. A *p*-value-based method, such as MaxP, can be conservative as it guards against the worst scenario and does not incorporate the non-null hypothesis. Lfdr incorporates the non-null density function of *p*-values and provides a different ranking of important SVGs. To account for the heterogeneity of *p*-value distributions under non-null from different studies, we use separate non-parametric estimations. STAREG does not involve any tuning parameters and is computationally scalable by combining the EM algorithm and PAVA. Moreover, although we focus on the replicability analysis of SVG detection from two SRT studies, STAREG can be easily extended to other modalities, such as scRNA-seq, ATAC-seq, and CITE-seq, among others.

A limitation of STAREG is that the current version of STAREG only considers the replicability analysis of two SRT studies. Replicability analysis of more than two studies requires the estimation of parameters that increase exponentially with the number of studies. We leave it for future research.

## 4 Methods

### 4.1 STAREG: model and algorithm

Suppose there are *m* genes in two SRT studies. We are interested in testing whether the *i*th gene shows any spatial expression pattern. The null hypothesis states that it does not exhibit any spatial expression pattern. STAREG aims to identify replicable SVGs by examining SRT data measured on two samples of the same tissue type. We first separately apply SVG detection method, such as SPARK (Sun *et al*., 2020) and/or SPARK-X (Zhu *et al*., 2021), to produce two sequences of well-calibrated *p*-values, and match them by genes. We assume *p*-values from the null follow the standard uniform distribution and *p*-values from the nonnull are stochastically smaller than the standard uniform distribution as smaller *p*-values indicate stronger evidence against the null (Cao *et al*., 2013). Denote paired *p*-values as (*p*_1_*_i_, p*_2_*_i_*)*, i* = 1*, . . ., m*. For *i*th gene, let *θ_ji_* be its hidden state in study *j* (*j* = 1, 2), where *θ_ji_* = 1 indicates *i*th gene is an SVG in study *j* and *θ_ji_* = 0 otherwise. We assume a two-group model for the two *p*-value sequences, respectively, where

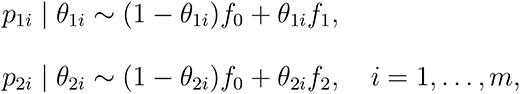

where *f*_0_ is the density function of *p*-values under the null, and *f*_1_ and *f*_2_ denote the non-null density functions for study 1 and study 2, respectively. The two studies share the same *p*-value distribution under the null. The heterogeneity across the two studies is accommodated through modeling density functions under the non-null separately by *f*_1_ and *f*_2_. Let *τ_i_* = (*θ*_1_*_i_, θ*_2_*_i_*)*, i* = 1*, . . ., m* denote the joint hidden states across two studies with prior probabilities *P* (*τ_i_* = (*k, l*)) = *ξ_kl_*, where *k, l* = 0, 1 and Σ*_k,l_ ξ_kl_* = 1, such that *τ_i_* ∈ {(0, 0), (0, 1), (1, 0), (1, 1)}. The replicability null hypothesis is

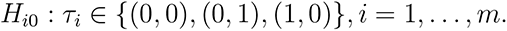

Lfdr is defined as the posterior probability of being null given data. We have

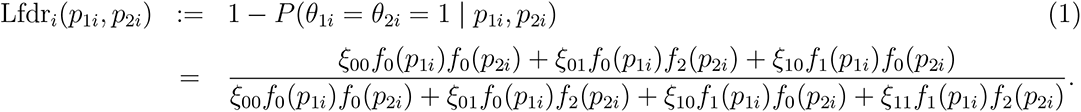

We assume the monotone likelihood ratio condition (Sun and Cai, 2007; Cao *et al*., 2013, 2022):

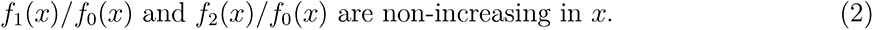

Let *≼* be the elementwise inequality in *R*^2^ (*x ≼ y* if and only if *x_k_ ≤ y_k_*for *k* = 1, 2). A function *g* : *R*^2^ *→ R* is monotone for this partial ordering if *x ≼ y* implies *g*(*x*) *≤ g*(*y*). From (2), we have that Lfdr*_i_*(*p*_1_*_i_, p*_2_*_i_*) is monotonically non-decreasing in (*p*_1_*_i_, p*_2_*_i_*). The rejection rule based on Lfdr*_i_* for testing replicability null *H_i_*_0_ is

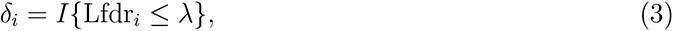

where *λ* is a threshold to be determined. We write total number of rejections as 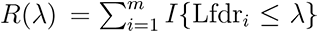, and number of false rejections as 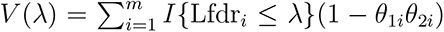

We have

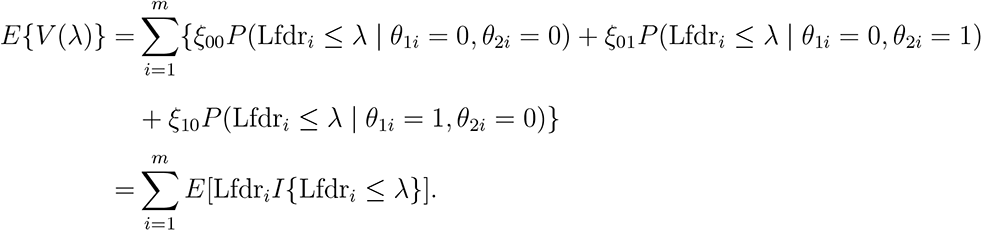

Write *a ∨ b* = max*{a, b}*. To control FDR of the replicability analysis, we need to find the critical value *λ* in (3). We estimate FDR by

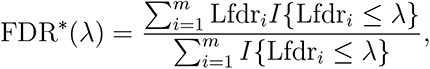

and define *λ_m_* = sup*{λ ∈* [0, 1] : FDR*^∗^*(*λ*) *≤ α*}. Reject *H_i_*_0_ if Lfdr*_i_ ≤ λ_m_*. This is the oracle case that assumes we know *ξ*_00_*, ξ*_01_*, ξ*_10_*, ξ*_11_*, f*_1_ and *f*_2_. We provide estimates of them in next section.

### 4.2 Estimates of unknowns and an adaptive procedure

Assume *f*_0_ follows a standard uniform distribution. Let 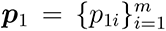 and 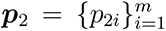 denote *p*-values from study 1 and study 2, respectively. Denote hidden states 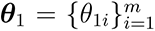 and 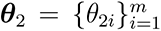. Under conditional independence of two *p*-value sequences given hidden states, the joint log likelihood function of (***p***_1_, ***p***_2_, ***θ***_1_, ***θ***_2_) is

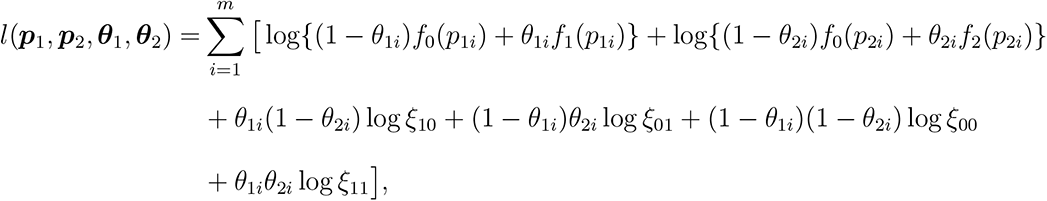

where hidden states ***θ***_1_ and ***θ***_2_ are latent variables. For scalable computation, we utilize EM algorithm (Dempster *et al*., 1977) in combination of pool-adjacent-violator-algorithm (PAVA) to efficiently estimate the unknowns (*ξ*_00_*, ξ*_01_*, ξ*_10_*, ξ*_11_*, f*_1_*, f*_2_) incorporating the monotone likelihood assumption (2) for *f*_1_ and *f*_2_ (see Supplementary Notes for details). *f*_1_ and *f*_2_ are estimated non-parametrically, which provides more flexibility. With the estimates (*ξ̂*_00_*, ξ̂*_01_*, ξ̂*_10_*, ξ̂*_11_*, f̂*_1_*, f̂*_2_), we obtain Lfdr estimates by:

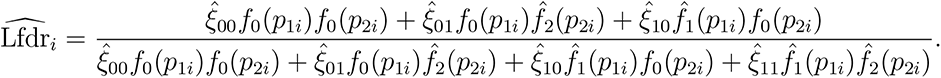

An estimate of *λ_m_*can be obtained through

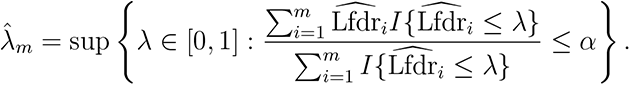

The replicability null hypothesis *H_i_*_0_ is rejected if 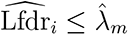, which means that gene *i* is identified as a replicable SVG across two SRT studies. This is equivalent to the step-up procedure

(Sun and Cai, 2007): let 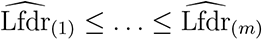 be the ordered statistics of 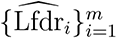 and denote by *H*_(1)_*, . . ., H*_(_*_m_*_)_ the corresponding ordered replicability null hypotheses, the procedure works as follows.

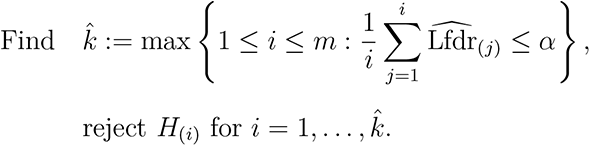

### 4.3 Comparison with other methods

BH procedure (Benjamini and Hochberg, 1995), a classic approach for FDR control in multiple testing, works as follows. Order *m p*-values from smallest to largest denoted as *p*_(1)_*, . . ., p*_(_*_m_*_)_. Find the largest *i*, denoted as *i*_0_, such that *p*_(_*_i_*_)_ is smaller than a threshold *αi/m.* Reject hypotheses corresponding to *p*_(1)_*, . . ., p*_(_*_i_* _)_. As proved in Benjamini and Hochberg (1995), this procedure controls the FDR at level *α* asymptotically when *p*-values under the null are independent and follow a standard uniform distribution. It is shown to be robust to positive dependence (Benjamini and Yekutieli, 2001).

To evaluate the performance of STAREG in identifying replicable SVGs from multiple SRT studies, we also implement the following two replicability analysis procedures for comparison.

- BH: an *ad hoc* approach that implements BH (Benjamini and Hochberg, 1995) procedure for each study and declares replicable findings as the intersection of significant findings from two studies.
- MaxP: a conservative method suggested in Benjamini et al. (2009) that applies the BH procedure (Benjamini and Hochberg, 1995) to the maximum *p*-values across two studies, max(*p*_1_*_i_, p*_2_*_i_*)*, i* = 1*, . . ., m*, for replicability analysis.

### 4.4 Data analysis details

#### 4.4.1 Mouse olfactory bulb data

The ST technology refers to the method developed in Ståhl *et al*. (2016), which positions individual tissue samples on arrays of spatially barcoded reverse transcription primers able to capture mRNA with the oligo(dT) tails. We obtained two replicates of ST data collected from mouse olfactory bulb published on the Spatial Research Website (https://www.spatialresearch.org/): Replicate 1 (file “MOB Replicate 1”) consists of 16, 573 genes measured on 267 spatial spots, and Replicate 8 (file “MOB Replicate 8”) contains 15, 288 genes measured on 234 spots. The two replicates were chosen because they are the least correlated of all 12 published ST data replicates. For each dataset, we filtered out genes that are expressed in less than 10% of the spatial locations and selected spatial locations with at least ten total read counts, resulting in 10, 373 genes on 265 spots in Replicate 1 dataset and 9, 671 genes on 232 spots in Replicate 8 dataset.

To validate the analysis results of the ST mouse olfactory bulb data, we downloaded and summarized three reference gene lists from the original ST study (Ståhl *et al*., 2016) and the Harmonizome database (Rouillard *et al*., 2016) (https://maayanlab.cloud/Harmonizome/). The first gene list from Ståhl *et al*. (2016) includes 10 genes with highly enriched expression in the mitral cell layer of mouse olfactory bulb: *Doc2g*, *Slc17a7*, *Reln*, *Cdhr1*, *Sv2b*, *Shisa3*, *Plcxd2*, *Nmb*, *Uchl1*, *Rcan2*. The second gene set obtained from the Allen Brain Atlas (Sunkin *et al*., 2013) includes 1, 894 genes with high or low expression in the main olfactory bulb relative to other tissues. The third gene set for validation contains 2, 031 genes differentially expressed in mouse olfactory bulb relative to other cell types and tissues from the BioGPS mouse cell type and tissue gene expression profiles dataset (Wu *et al*., 2013).

#### 4.4.2 Human breast cancer data

The human breast cancer data measured with ST was downloaded from the Spatial Research Website (https://www.spatialresearch.org/) with file “Breast Cancer Layer 2”, consisting of 14, 789 genes on 251 spatial spots. The 10X Visium platform builds upon original ST technology with improved resolution. We downloaded another human breast cancer data from 10X Visium spatial gene expression repository (https://www.10xgenomics.com/resources/datasets/) for replicability analysis, including 36, 601 genes measured on 3, 798 spatial locations. After filtering out genes expressed in less than 10% of the locations and spatial locations with less than ten total read counts, we obtained 5, 262 genes on 250 spots in the ST dataset and 11, 993 genes on 3, 798 locations in the 10X Visium dataset.

We provided two lists of reference gene sets related to human breast cancer from previous literature and publicly available database to validate replicable SVGs identified by different methods. The first list of 3, 505 human breast cancer-related genes was summarized from six different datasets collected in the Harmonizome database (Rouillard *et al*., 2016): OMIM gene-disease associations (Amberger et al., 2015), PhosphoSitePlus Phosphosite-disease associations (Hornbeck et al., 2015), DISEASES text-mining gene-disease association evidence scores (Pletscher-Frankild et al., 2015), GAD gene-disease associations (Becker et al., 2004), GWAS catalog SNP-phenotype associations (Welter et al., 2014), and GWASdb SNP-disease associations (Li et al., 2012). The second reference gene list includes 4, 842 genes differentially expressed between cell-type clusters in human breast cancers (Wu *et al*., 2021).

#### 4.4.3 Mouse cerebellum data

Slide-seq is a technology that provides scalable methods for obtaining SRT data at resolutions comparable to single cells (Rodriques *et al*., 2019). Slide-seqV2 is built on Slide-seq, combining improvements in library generation, bead synthesis, and array indexing to achieve a ten-fold higher RNA capture efficiency than Slide-seq (Stickels *et al*., 2021). We obtained two SRT datasets of mouse cerebellum measured with Slide-seq and Slide-seqV2 from Broad Institute’s single-cell repository (https://singlecell.broadinstitute.org/single_cell), with IDs SCP354 and SCP948, respectively. For the Slide-seq data, we used the dataset summarized in the file “Puck_180819_12”, which contains 19, 782 genes measured on 32, 701 beads. The Slide-seqV2 data contains 23, 096 genes on 39, 496 beads. For the Slide-seq dataset, we first filtered out beads that were not assigned clusters in the original study (Rodriques *et al*., 2019). For the Slide-seqV2 dataset, we cropped regions of interest by filtering out beads with UMIs less than 100 following Cable *et al*. (2022). Mitochondrial genes and genes that were not expressed on any location were filtered out from the two datasets, and beads with zero total expression counts were removed, resulting in 18, 082 genes on 28, 352 beads in the Slide-seq dataset and 20, 117 genes on 11, 626 beads in the Slide-seqV2 dataset for replicability analysis. We downloaded two reference gene lists from previously published literature to provide unbiased validations for the results of our replicability analysis. First, we obtained a list of highly differentially expressed genes across all cell clusters in mouse cerebellar cortex from Kozareva et al. (2021). Let log FC denote logarithmic fold changes. By filtering out genes with | log FC| < 1.5, we used a final set of 995 marker genes for the validation. The second validation set was obtained from the Harmonizome database (Rouillard *et al*., 2016), including 2, 431 genes with high or low expression in cerebellum relative to other cell types and tissues from the BioGPS mouse cell type and tissue gene expression profiles dataset (Wu *et al*., 2013).

#### 4.4.4 Computational time

We evaluated the computational efficiency of different methods in the replicability analysis of three pairs of SRT datasets: mouse olfactory bulb data, human breast cancer data, and mouse cerebellum data. Table 2 summarizes the data information and computational time. Computations were carried out in an Intel(R) Core(TM) i7-9750H 2.6GHz CPU with 64.0 GB RAM laptop. We see that all methods are fast, while STAREG takes longer time than BH and MaxP. Such differences are ignorable for practical use.

**Table 2:** Computational time (in seconds) for replicability analysis in different datasets: mouse olfactory bulb (MOB), human breast cancer (HBC), and mouse cerebellum (MC).

| Dataset | # of genes/samples |  | # of genes common to both studies | BH | MaxP | STAREG |
| --- | --- | --- | --- | --- | --- | --- |
|  | study 1 | study 2 |  |  |  |  |
| MOB | 10,373/265 | 9,671/232 | 9,329 | 0.005 | 0.026 | 2.635 |
| HBC | 5,262/250 | 11,993/3,798 | 4,943 | 0.004 | 0.015 | 0.157 |
| MC | 18,082/28,352 | 20,117/11,626 | 16,873 | 0.006 | 0.047 | 4.301 |

## Data availability

This study analyzed three pairs of SRT datasets. All data are publicly available and can be acquired from corresponding websites. Two replicates of the ST data from mouse olfactory bulb can be downloaded from the Spatial Research Website at https://www.spatialresearch.org/resources-published-datasets/doi-10-1126science-aaf2403 (files “MOB Replicate 1” and “MOB Replicate 8”). The ST data from human breast cancer is presented in the file “Breast Cancer Layer 2” at the Spatial Research Website. The 10X Visium data from human breast cancer is available in the 10X Visium spatial gene expression repository under item Visium Spatial Gene Expression 1.1.0 (https://www.10xgenomics.com/resources/datasets/human-breast-cancer-block-a-section-1-1-standard-1-1-0). The Slide-seq data and Slide-seqV2 data from the mouse cerebellum are available at the Broad Institute’s single-cell repository (https://singlecell.broadinstitute.org/single_cell/) under IDs SCP354 (file “Puck_180819_12”) and SCP948, respectively.

## Code availability

We implement the STAREG method in an open-source R package, STAREG, which is freely available at https://github.com/hongyuan-cao/STAREG. The R source code to reproduce our analysis can be found at https://github.com/hongyuan-cao/STAREG-Analysis.

## Acknowledgements

This work was partially supported by the Postdoctoral Research Foundation of China (No. 801212021410) to Yan Li. We thank Joel E. Cohen for correcting some typos.

## Author information

### Contributions

H.C. conceived the main idea. Y.L. performed the simulations and real data analysis. All authors contributed to the methodology and data analysis discussions. Y.L. and H.C. wrote the manuscript. X.Z. and H.C. revised the manuscript.

