## Supplementary Material for "STAREG: an empirical Bayesian approach to detect replicable spatially variable genes in spatial transcriptomic studies"

[1]School of Mathematics, Jilin University, Changchun, Jilin 130012, China. [2]Department of Biostatistics, University of Michigan, Ann Arbor, MI 48109, USA. [3]Department of Molecular and Human Genetics, Baylor College of Medicine, Houston, TX 77030, USA. [4]Department of Statistics, Texas A&M University, College Station, TX 77843, USA. [5]Department of Statistics, Florida State University, Tallahassee, FL 32306, USA.

### A Estimate unknowns with the EM algorithm

In this section, we show how to estimate unknown parameters and functions involved in $\text{Lfdr}_i$ from available paired $p$-value sequences: $(p_{1i}, p_{2i}), i = 1, \ldots, m$. Based on the four-group model, the posterior probability mass function $P(\theta_{1i}, \theta_{2i} | p_{1i}, p_{2i})$ can be written as

$$
\begin{aligned}
\gamma_{i,00} &= P(\theta_{1i} = 0, \theta_{2i} = 0 \mid p_{1i}, p_{2i}) \\
&= \frac{\xi_{00} f_0(p_{1i}) f_0(p_{2i})}{\xi_{00} f_0(p_{1i}) f_0(p_{2i}) + \xi_{01} f_0(p_{1i}) f_2(p_{2i}) + \xi_{10} f_1(p_{1i}) f_0(p_{2i}) + \xi_{11} f_1(p_{1i}) f_2(p_{2i})}, \\
\gamma_{i,01} &= P(\theta_{1i} = 0, \theta_{2i} = 1 \mid p_{1i}, p_{2i}) \\
&= \frac{\xi_{01} f_0(p_{1i}) f_2(p_{2i})}{\xi_{00} f_0(p_{1i}) f_0(p_{2i}) + \xi_{01} f_0(p_{1i}) f_2(p_{2i}) + \xi_{10} f_1(p_{1i}) f_0(p_{2i}) + \xi_{11} f_1(p_{1i}) f_2(p_{2i})}, \\
\gamma_{i,10} &= P(\theta_{1i} = 1, \theta_{2i} = 0 \mid p_{1i}, p_{2i}) \\
&= \frac{\xi_{10} f_1(p_{1i}) f_0(p_{2i})}{\xi_{00} f_0(p_{1i}) f_0(p_{2i}) + \xi_{01} f_0(p_{1i}) f_2(p_{2i}) + \xi_{10} f_1(p_{1i}) f_0(p_{2i}) + \xi_{11} f_1(p_{1i}) f_2(p_{2i})}, \\
\gamma_{i,11} &= P(\theta_{1i} = 1, \theta_{2i} = 1 \mid p_{1i}, p_{2i}) \\
&= \frac{\xi_{11} f_1(p_{1i}) f_2(p_{2i})}{\xi_{00} f_0(p_{1i}) f_0(p_{2i}) + \xi_{01} f_0(p_{1i}) f_2(p_{2i}) + \xi_{10} f_1(p_{1i}) f_0(p_{2i}) + \xi_{11} f_1(p_{1i}) f_2(p_{2i})},
\end{aligned}
$$

where $f_0$ is the density function of standard uniform distribution. We need to estimate unknown parameters $\xi_{00}, \xi_{01}, \xi_{10}, \xi_{11}$ and unknown functions $f_1, f_2$. With a reasonable initialization of the unknowns $(\hat{\xi}_{00}^{(0)}, \hat{\xi}_{01}^{(0)}, \hat{\xi}_{10}^{(0)}, \hat{\xi}_{11}^{(0)}, \hat{f}_1^{(0)}, \hat{f}_2^{(0)})$, we can derive an EM algorithm (Dempster *et al.*, 1977) by iteratively implementing the following two steps.

**E-step:** Given current estimates of $(\hat{\xi}_{00}^{(t)}, \hat{\xi}_{01}^{(t)}, \hat{\xi}_{10}^{(t)}, \hat{\xi}_{11}^{(t)}, \hat{f}_1^{(t)}, \hat{f}_2^{(t)})$, calculate $\gamma_{i,00}^{(t)}, \gamma_{i,01}^{(t)}, \gamma_{i,10}^{(t)}, \gamma_{i,11}^{(t)}$, respectively. Define conditional expectation of the log-likelihood function as

$$
\begin{aligned}
&D(\xi_{00}, \xi_{01}, \xi_{10}, \xi_{11}, f_1, f_2 \mid \hat{\xi}_{00}^{(t)}, \hat{\xi}_{01}^{(t)}, \hat{\xi}_{10}^{(t)}, \hat{\xi}_{11}^{(t)}, \hat{f}_1^{(t)}, \hat{f}_2^{(t)}) \\
=&\mathbb{E}_{\boldsymbol{\theta}_1, \boldsymbol{\theta}_2 | \hat{\xi}_{00}^{(t)}, \hat{\xi}_{01}^{(t)}, \hat{\xi}_{10}^{(t)}, \hat{\xi}_{11}^{(t)}, \hat{f}_1^{(t)}, \hat{f}_2^{(t)}} [l(\boldsymbol{p}_1, \boldsymbol{p}_2, \boldsymbol{\theta}_1, \boldsymbol{\theta}_2)] \\
=&\sum_{i=1}^{m} [(\gamma_{i,00}^{(t)} + \gamma_{i,01}^{(t)}) \log f_0(p_{1i}) + (\gamma_{i,10}^{(t)} + \gamma_{i,11}^{(t)}) \log f_1(p_{1i}) + (\gamma_{i,00}^{(t)} + \gamma_{i,10}^{(t)}) \log f_0(p_{2i}) \\
&+ (\gamma_{i,01}^{(t)} + \gamma_{i,11}^{(t)}) \log f_2(p_{2i})] + \sum_{i=1}^{m} [\gamma_{i,00}^{(t)} \log \xi_{00} + \gamma_{i,01}^{(t)} \log \xi_{01} + \gamma_{i,10}^{(t)} \log \xi_{10} + \gamma_{i,11}^{(t)} \log \xi_{11}].
\end{aligned}
$$

**M-step:** Update $(\hat{\xi}_{00}^{(t+1)}, \hat{\xi}_{01}^{(t+1)}, \hat{\xi}_{10}^{(t+1)}, \hat{\xi}_{11}^{(t+1)}, \hat{f}_1^{(t+1)}, \hat{f}_2^{(t+1)})$ by maximizing the conditional expectation of the log-likelihood function $D(\xi_{00}, \xi_{01}, \xi_{10}, \xi_{11}, f_1, f_2 \mid \hat{\xi}_{00}^{(t)}, \hat{\xi}_{01}^{(t)}, \hat{\xi}_{10}^{(t)}, \hat{\xi}_{11}^{(t)}, \hat{f}_1^{(t)}, \hat{f}_2^{(t)})$ subject to constraint $\xi_{00} + \xi_{01} + \xi_{10} + \xi_{11} = 1$. We obtain

$$
\begin{aligned}
\hat{\xi}_{00}^{(t+1)} &= \frac{\sum_{i=1}^{m} \gamma_{i,00}}{m}, \\
\hat{\xi}_{01}^{(t+1)} &= \frac{\sum_{i=1}^{m} \gamma_{i,01}}{m}, \\
\hat{\xi}_{10}^{(t+1)} &= \frac{\sum_{i=1}^{m} \gamma_{i,10}}{m}, \\
\hat{\xi}_{11}^{(t+1)} &= \frac{\sum_{i=1}^{m} \gamma_{i,11}}{m},
\end{aligned}
$$

and

$$
\hat{f}_1^{(t+1)} = \arg\max_{\tilde{f}_1 \in \mathbb{H}} \left\{ \sum_{i=1}^{m} (\gamma_{i,10}^{(t)} + \gamma_{i,11}^{(t)}) \log \tilde{f}_1(p_{1i}) \right\}, \tag{S1}
$$

$$
\hat{f}_2^{(t+1)} = \arg\max_{\tilde{f}_2 \in \mathbb{H}} \left\{ \sum_{i=1}^{m} (\gamma_{i,01}^{(t)} + \gamma_{i,11}^{(t)}) \log \tilde{f}_2(p_{2i}) \right\}, \tag{S2}
$$

where $\mathbb{H}$ is the set of $p$-value density functions under the non-null. We repeat the above

**E-step** and **M-step** until the algorithm converges.

Next we provide specific steps to solve (S1) and (S2) using the Pool-Adjacent-Violators Algorithm (PAVA) (Robertson *et al.*, 1988) under the monotone likelihood ratio assumption (Sun and Cai, 2007; Cao *et al.*, 2013, 2022). Denote $Q_{1i}^{(t)} = \gamma_{i,10}^{(t)} + \gamma_{i,11}^{(t)}$ and $Q_{2i}^{(t)} = \gamma_{i,01}^{(t)} + \gamma_{i,11}^{(t)}, i = 1, \ldots, m$. Let $0 = p_{1(0)} \leq p_{1(1)} \leq \cdots \leq p_{1(m)}$ be the order statistics of $\boldsymbol{p}_1$ and denote $Q_{1(i)}^{(t)}$ as corresponding $Q_{1i}^{(t)}$ associated with $p_{1(i)}$. Let $0 = p_{2(0)} \leq p_{2(1)} \leq \cdots \leq p_{2(m)}$ be the order statistics of $\boldsymbol{p}_2$ and denote $Q_{2(i)}^{(t)}$ as corresponding $Q_{2i}^{(t)}$ associated with $p_{2(i)}$. Define $y_{1i} = f_1\left(p_{1(i)}\right)$ and $y_{2i} = f_2\left(p_{2(i)}\right)$, we can write (S1) and (S2) as

$$\hat{f}_1^{(t+1)} = \arg\max_{y_{1i}\in\mathcal{M}_1} \left\{\sum_{i=1}^m Q_{1(i)}^{(t)} \log y_{1i}\right\}, \text{ subject to } \sum_{i=1}^m y_{1i}(p_{1(i)} - p_{1(i-1)}) = 1,$$
$$\hat{f}_2^{(t+1)} = \arg\max_{y_{2i}\in\mathcal{M}_2} \left\{\sum_{i=1}^m Q_{2(i)}^{(t)} \log y_{2i}\right\}, \text{ subject to } \sum_{i=1}^m y_{2i}(p_{2(i)} - p_{2(i-1)}) = 1,$$

where $\mathcal{M}_1 = \{(y_{11}, \ldots, y_{1m}) \mid y_{11} \geq \cdots \geq y_{1m} \geq 0\}$ and $\mathcal{M}_2 = \{(y_{21}, \ldots, y_{2m}) \mid y_{21} \geq \cdots \geq y_{2m} \geq 0\}$.

By the Lagrangian multiplier, the objective functions we want to maximize are

$$\sum_{i=1}^m Q_{1(i)}^{(t)} \log y_{1i} + \lambda_1 \left\{\sum_{i=1}^m y_{1i}(p_{1(i)} - p_{1(i-1)}) - 1\right\},$$
$$\sum_{i=1}^m Q_{2(i)}^{(t)} \log y_{2i} + \lambda_2 \left\{\sum_{i=1}^m y_{2i}(p_{2(i)} - p_{2(i-1)}) - 1\right\}.$$

Taking derivatives with respect to $y_{1i}, \lambda_1$ and $y_{2i}, \lambda_2$, respectively, we have

$$\hat{\lambda}_1 = -\sum_{i=1}^m Q_{1(i)}^{(t)},\ \hat{y}_{1i} = \frac{Q_{1(i)}^{(t)}}{Q_1^{(t)}(p_{1(i)} - p_{1(i-1)})},$$
$$\hat{\lambda}_2 = -\sum_{i=1}^m Q_{2(i)}^{(t)},\ \hat{y}_{2i} = \frac{Q_{2(i)}^{(t)}}{Q_2^{(t)}(p_{2(i)} - p_{2(i-1)})},$$

where $Q_1^{(t)} = \sum_{i=1}^m Q_{1i}^{(t)}$ and $Q_2^{(t)} = \sum_{i=1}^m Q_{2i}^{(t)}$.

To incorporate the monotone constraints on $y_{1i}$ and $y_{2i}$, we minimize

$$\sum_{i=1}^m \left\{ -Q_{1(i)}^{(t)} \log y_{1i} + Q_1^{(t)}(p_{1(i)} - p_{1(i-1)}) y_{1i} \right\} = \sum_{i=1}^m Q_{1(i)}^{(t)} \left\{ -\log y_{1i} - \frac{-Q_1^{(t)} \left( p_{1(i)} - p_{1(i-1)} \right)}{Q_{1(i)}^{(t)}} y_{1i} \right\},$$

subject to $y_{11} \geq \cdots \geq y_{1m}$, and minimize

$$\sum_{i=1}^m \left\{ -Q_{2(i)}^{(t)} \log y_{2i} + Q_2^{(t)}(p_{2(i)} - p_{2(i-1)}) y_{2i} \right\} = \sum_{i=1}^m Q_{2(i)}^{(t)} \left\{ -\log y_{2i} - \frac{-Q_2^{(t)} \left( p_{2(i)} - p_{2(i-1)} \right)}{Q_{2(i)}^{(t)}} y_{2i} \right\},$$

subject to $y_{21} \geq \cdots \geq y_{2m}$.

Let

$$(\hat{u}_{11}, \ldots, \hat{u}_{1m}) = \arg \min_{u_{11}, \ldots, u_{1m}} \sum_{i=1}^m Q_{1(i)}^{(t)} \left( u_{1i} - \frac{-Q_1^{(t)} \left( p_{1(i)} - p_{1(i-1)} \right)}{Q_{1(i)}^{(t)}} \right)^2$$

subject to $u_{11} \geq u_{12} \geq \cdots \geq u_{1m}$, and

$$(\hat{u}_{21}, \ldots, \hat{u}_{2m}) = \arg \min_{u_{21}, \ldots, u_{2m}} \sum_{i=1}^m Q_{2(i)}^{(t)} \left( u_{2i} - \frac{-Q_2^{(t)} \left( p_{2(i)} - p_{2(i-1)} \right)}{Q_{2(i)}^{(t)}} \right)^2$$

subject to $u_{21} \geq u_{22} \geq \cdots \geq u_{2m}$. The solutions take the max-min form

$$\hat{u}_{1i} = \max_{b \geq i} \min_{a \leq i} \frac{-Q_1^{(t)} \sum_{k=a}^b \left( p_{1(k)} - p_{1(k-1)} \right)}{\sum_{k=a}^b Q_{1(k)}^{(t)}},$$
$$\hat{u}_{2i} = \max_{b \geq i} \min_{a \leq i} \frac{-Q_2^{(t)} \sum_{k=a}^b \left( p_{2(k)} - p_{2(k-1)} \right)}{\sum_{k=a}^b Q_{2(k)}^{(t)}},$$

which can be obtained by PAVA (Robertson *et al.*, 1988). Our final estimates are given by $\hat{y}_{1i} = -\frac{1}{\hat{u}_{1i}}$ and $\hat{y}_{2i} = -\frac{1}{\hat{u}_{2i}}$ for $i = 1, \ldots, m$ according to Theorem 3.1 of Barlow and Brunk (1972).

### B Simulation studies

Simulations were performed to evaluate FDR control and the power of different methods. Let $h_i$ indicate joint states for $m$ replicability null hypotheses $H_{i0} : (\theta_{1i}, \theta_{2i}) \in \{(0,0), (0,1), (1,0)\}, i = 1, \ldots, m$, such that

$$h_i = \begin{cases} 0, & \text{if } (\theta_{1i}, \theta_{2i}) = (0,0), \\ 1, & \text{if } (\theta_{1i}, \theta_{2i}) = (0,1), \\ 2, & \text{if } (\theta_{1i}, \theta_{2i}) = (1,0), \\ 3, & \text{if } (\theta_{1i}, \theta_{2i}) = (1,1). \end{cases}$$

We generated $h_i$ for $i = 1, \ldots, m$ from a multinomial distribution with prior probabilities:

$$\mathbb{P}(h_i = 0) = \xi_{00},$$
$$\mathbb{P}(h_i = 1) = \xi_{01},$$
$$\mathbb{P}(h_i = 2) = \xi_{10},$$
$$\mathbb{P}(h_i = 3) = \xi_{11}.$$

Then we simulated separate hidden states for study $j$ ($j = 1, 2$), $\boldsymbol{\theta}_j = \{\theta_{ji}\}_{i=1}^m$ from $h_i, i = 1, \ldots, m$.

#### B.1 Simulation study 1

We first performed simulation studies based on normal distributions and $z$-tests. Denote $N(\mu, \sigma^2)$ a normal distribution with mean $\mu$ and variance $\sigma^2$. For study $j$, we independently generated $z$-statistics from $X_{ji} \sim N(0, \sigma_j^2)$ if $\theta_{ji} = 0$, and from $X_{ji} \sim N(\mu_j, \sigma_j^2)$ if $\theta_{ji} = 1$, where $\mu_j > 0$. One-sided $p$-value for each gene was calculated by $p_{ji} = 1 - \Phi(X_{ji}/\sigma_j), j = 1, 2; i = 1, \ldots, m$, where $\Phi(\cdot)$ is the cumulative distribution function of standard normal distribution $N(0,1)$.

Set $m = 10,000$, $\xi_{11} = 0.05$ and let $\xi_{01} = \xi_{10}$ take values from 0.04 to 0.2. Corresponding $\xi_{00}$ can be calculated by $\xi_{00} = 1-\xi_{01}-\xi_{10}-\xi_{11}$. At a target FDR level of 0.05, FDR and power of different methods were calculated from 100 replications in each setting. Fig. S1a shows the empirical FDR and power over a range of values for $\xi_{01}$ under different non-null settings. We see that MaxP was overly conservative in all settings. BH failed to control FDR when the probability of inconsistency between two studies is relatively large (e.g., $\xi_{01} = \xi_{10} > 0.14$). STAREG properly controlled FDR in all settings and showed higher power when the FDR of BH is under control. Because BH failed to control FDR, we also plot the power of different methods with corresponding empirical FDR obtained from one replication under the setting of $\xi_{00} = 0.95, \xi_{01} = \xi_{10} = 0.025$ and $\xi_{11} = 0.05$ (Fig. S1b) to have a fair comparison. We observe that STAREG showed higher power than the other methods at the same empirical FDR level.

#### B.2 Simulation study 2

We performed realistic simulations based on Replicate 1 and Replicate 8 of the ST data from mouse olfactory bulb with parameters inferred from SPARK (Sun *et al.*, 2020) (details of the studies can be found in the main Results section for analyzing mouse olfactory bulb data). In all realistic simulations, we used pre-specified $\xi_{00}, \xi_{01}, \xi_{10}$ and $\xi_{11}$ to generate hidden states $\boldsymbol{\theta}_1$ and $\boldsymbol{\theta}_2$ for $m$ genes. We then separately generated count data in two studies following the simulation strategy in Sun *et al.* (2020) based on $\boldsymbol{\theta}_1$ and $\boldsymbol{\theta}_2$. Specifically, in study $j$ ($j = 1, 2$), for each gene, in turn, the read counts on the spot $i$ ($i = 1, \ldots, n$, $n = 265$ for study 1 based on Replicate 1 and $n = 232$ for study 2 based on Replicate 8) were simulated from the following model:

$$y_i \sim \text{Poisson}(N_i \lambda_i), \ \log \lambda_i = \beta_i + \epsilon_i,$$

where $N_i$ denotes total read counts of all genes on the spot $i$, which can be obtained from data; $\lambda_i$ is the unknown relative expression level of the focal gene; the intercept $\beta_i$ represents the mean value of $\log \lambda_i$; and the error term $\epsilon_i \sim N(0, \tau_j^2)$ measures random noise independent of spatial locations, with the same variance across all spatial spots in the same study. The standard deviation $\tau_j$ for study $j$ was specified as $0.2, 0.5$, or $0.8$ in the simulations. For non-SVGs, we set the intercept $\beta_i$ to be constant across all spots and equal to the median of the intercepts estimated by SPARK in corresponding mouse olfactory bulb replicates ($\beta_i = -10.46$ for study 1 and $\beta_i = -9.98$ for study 2). For SVGs, we introduced the spatial expression patterns by dividing the $n$ spots into two groups based on the three main patterns in mouse olfactory bulb as illustrated in Fig. 2a. In the low expression group, $\beta_i$ is set to be $-10.46$ for study 1 and $-9.98$ for study 2; and in the high expression group, $\beta_i$ is set to be two-fold (weak signal), three-fold (moderate signal) or four-fold (strong signal) higher than that in the low expression group on rate parameter scale, e.g., $e^{\beta_i} = 2 \cdot e^{-10.46}$ means $\beta_i$ is two-fold higher than $-10.46$. Finally, $\lambda_i$ can be simulated through $\lambda_i = e^{\beta_i + \epsilon_i}$, and $y_i$ was generated from the Poisson distribution with parameter $N_i \lambda_i$.

Based on the above realistic simulation strategy, we varied the value of $\tau_j$ and signal strengths to produce paired count data for $m = 10,000$ genes. Then we used SPARK (Sun *et al.*, 2020) to produce well-calibrated $p$-values for each study, and performed replicability analysis of paired $p$-value sequences to evaluate FDR control (Fig. S2) and power (Fig. S3) of different methods. To have a fair comparison, we also calculated the power of different methods with corresponding empirical FDR based on a range of FDR cutoffs under the setting of $\xi_{00} = 0.9, \xi_{01} = \xi_{10} = 0.025$, and $\xi_{11} = 0.05$. Fig. 1b shows the results of simulations based on three patterns under the setting of $\tau_1 = \tau_2 = 0.2$ with a strong signal for study 1 and a weak signal for study 2. Additional simulation results with different noise levels ($\tau_j$) and signal strengths can be found in Fig. S4. Overall, STAREG controlled FDR at a pre-specified significance level and has higher power than competing methods. Moreover, it is interesting to note that all methods produced higher power in Pattern II- and Pattern III-based simulations than in Pattern I-based simulations, which is probably due to the clearer spatial expression patterns.

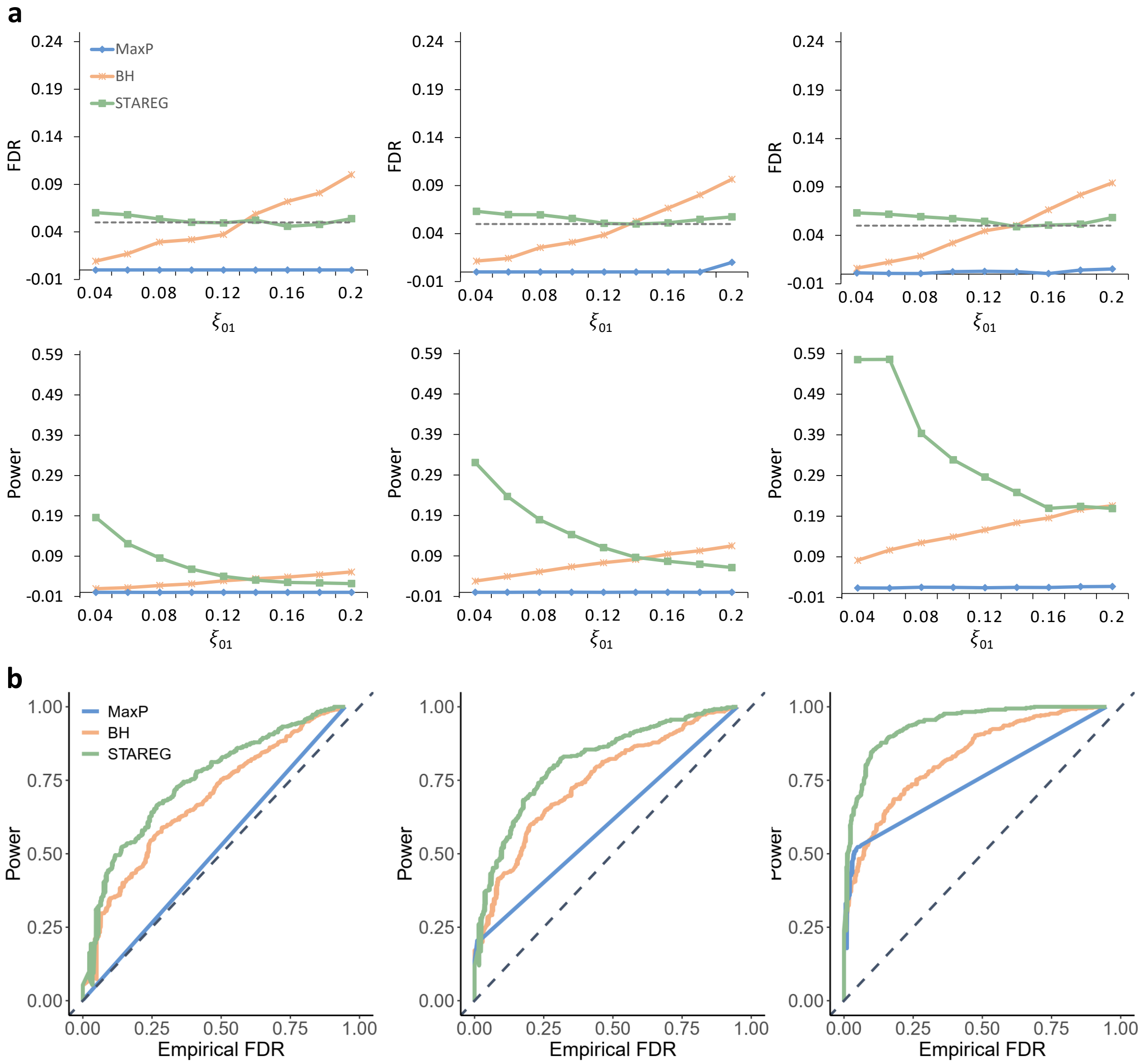


Figure S1: Performance comparisons of different methods in simulation study 1.
(**a**) Plots of empirical FDR (first row) and power (second row) over different values of $\xi_{01}$ (x-axis). Simulations were conducted under the setting of $m = 10,000$, $\xi_{11} = 0.05$ and $\xi_{01} = \xi_{10}$. The target FDR level is 0.05 (horizontal dashed line in the FDR plots), and the results were calculated over 100 replications. Each column corresponds to a different non-null setting (left: $\mu_1 = \mu_2 = 2, \sigma_1 = \sigma_2 = 1$; middle: $\mu_1 = 2, \mu_2 = 2.5, \sigma_1 = \sigma_2 = 1$; right: $\mu_1 = \mu_2 = 2, \sigma_1 = 1, \sigma_2 = 0.5$).
(**b**) Plots of power (y-axis) over corresponding empirical FDR (x-axis) at a range of FDR cutoffs. Simulations were performed with $m = 10,000, \xi_{00} = 0.9, \xi_{01} = \xi_{10} = 0.025$ and $\xi_{11} = 0.05$ under different non-null settings (left: $\mu_1 = \mu_2 = 2, \sigma_1 = \sigma_2 = 1$; middle: $\mu_1 = 2, \mu_2 = 2.5, \sigma_1 = \sigma_2 = 1$; right: $\mu_1 = \mu_2 = 2, \sigma_1 = 1, \sigma_2 = 0.5$). The diagonal dashed line with slope 1 is used as a reference.

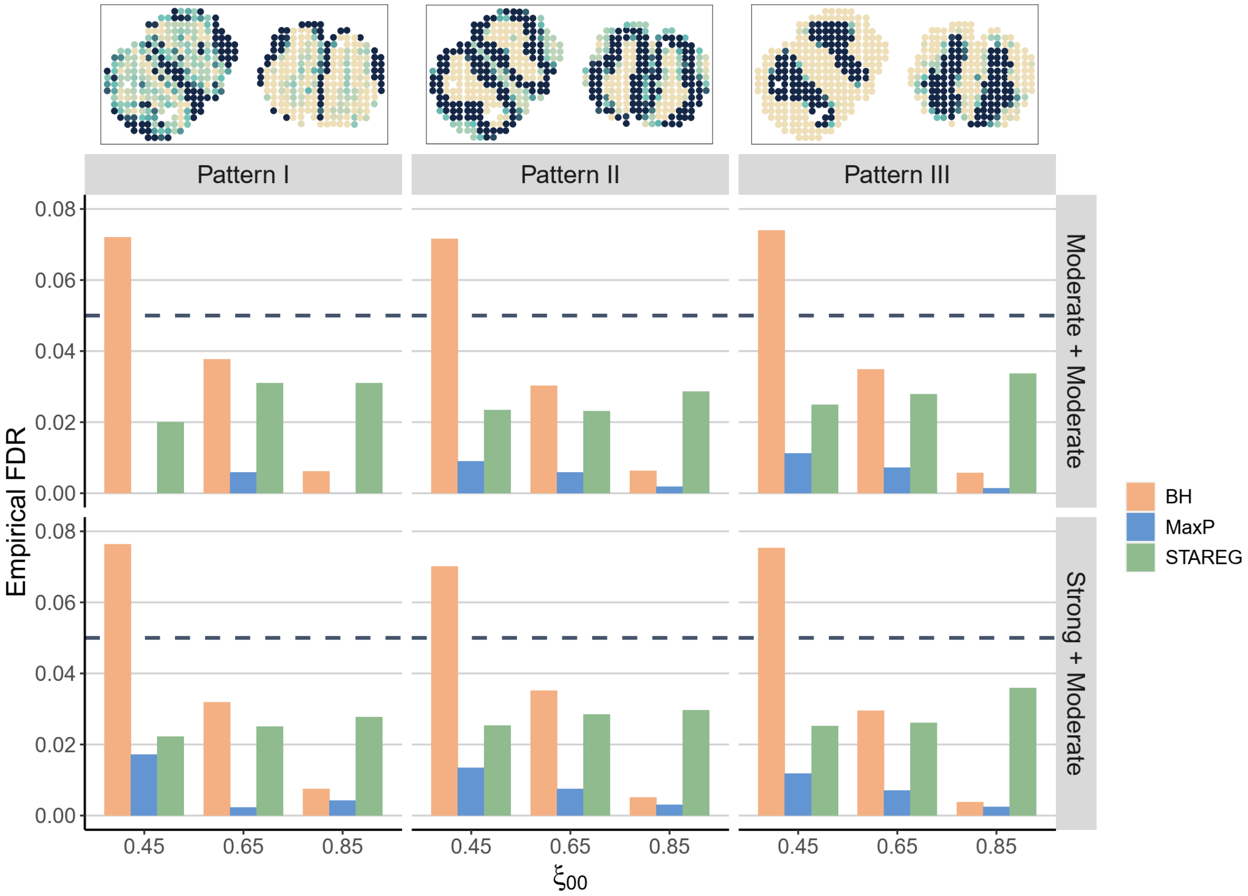


Figure S2: FDR control of different methods in simulation study 2. Simulations were performed for $m = 10,000$ genes and the target FDR level is 0.05. In each simulation, two SRT datasets were independently generated from corresponding mouse olfactory bulb data with the same noise level ($\tau_1 = \tau_2 = 0.2$) and different prior probabilities (left: $\xi_{00} = 0.85, \xi_{01} = \xi_{10} = 0.05, \xi_{11} = 0.05$; middle: $\xi_{00} = 0.65, \xi_{01} = \xi_{10} = 0.15, \xi_{11} = 0.05$; right: $\xi_{00} = 0.45, \xi_{01} = \xi_{10} = 0.25, \xi_{11} = 0.05$). Each column corresponds to a different spatial expression pattern, as illustrated at the top (left to right: Pattern I-III). Each row corresponds to a different signal strength setting for the two studies (top: moderate signal for both studies; bottom: strong signal for study 1 and moderate signal for study 2). The empirical FDR was averaged across ten replications. The horizontal dashed line represents the target FDR level of 0.05.

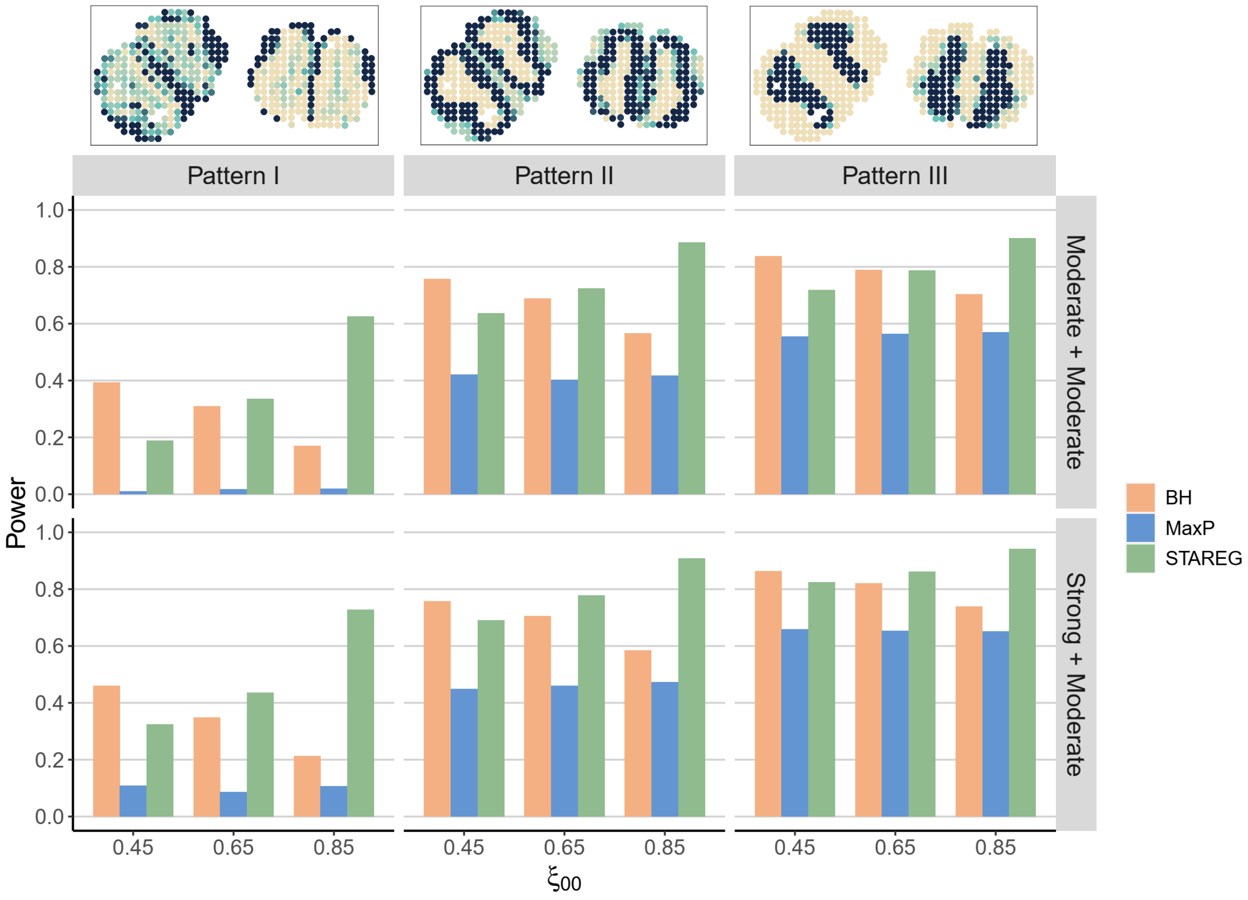


Figure S3: Power comparison of different methods in simulation study 2. Simulations were performed for $m = 10,000$ genes and the target FDR level is 0.05. In each simulation, two SRT datasets were independently generated from corresponding mouse olfactory bulb data with the same noise level ($\tau_1 = \tau_2 = 0.2$) and different prior probabilities (left: $\xi_{00} = 0.85, \xi_{01} = \xi_{10} = 0.05, \xi_{11} = 0.05$; middle: $\xi_{00} = 0.65, \xi_{01} = \xi_{10} = 0.15, \xi_{11} = 0.05$; right: $\xi_{00} = 0.45, \xi_{01} = \xi_{10} = 0.25, \xi_{11} = 0.05$). Each column corresponds to a different spatial expression pattern, as illustrated at the top (left to right: Pattern I-III). Each row corresponds to a different signal strength setting for the two studies (top: moderate signal for both studies; bottom: strong signal for study 1 and moderate signal for study 2). The empirical FDR was averaged across ten replications.

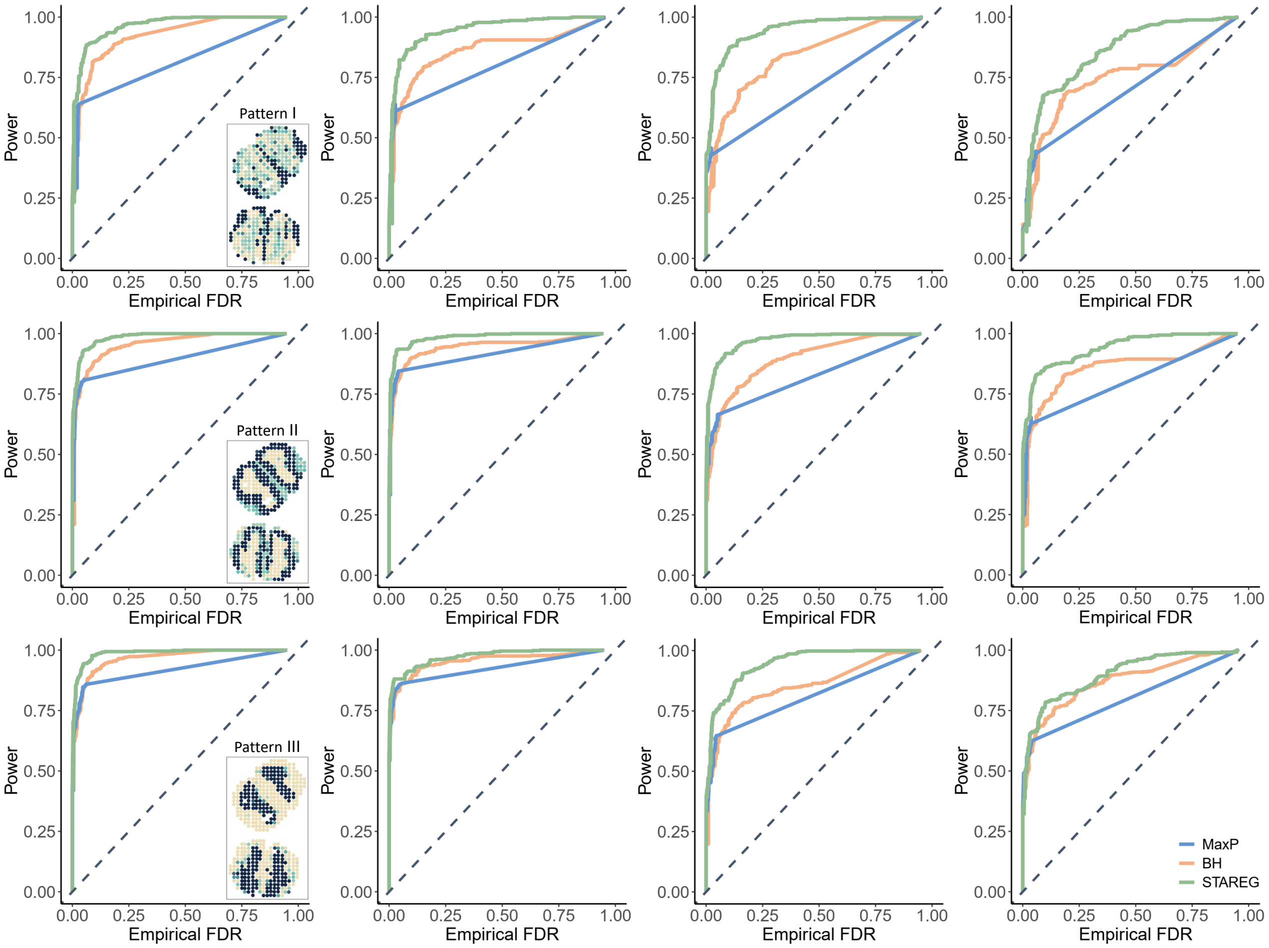


Figure S4: Plots of power (y-axis) over corresponding empirical FDR (x-axis) of different methods at a range of FDR cutoffs in simulation study 2. Simulations were performed under the setting of $m = 10,000$, $\xi_{00} = 0.9, \xi_{01} = \xi_{10} = 0.025$ and $\xi_{11} = 0.05$. The signal strengths were set to be strong for study 1 and moderate for study 2 across all simulations. In each setting, two SRT datasets were independently generated from corresponding mouse olfactory bulb replicates with different noise levels (left to right: $\tau_1 = 0.2, \tau_2 = 0.5$; $\tau_1 = \tau_2 = 0.5$; $\tau_1 = 0.5, \tau_2 = 0.8$; $\tau_1 = \tau_2 = 0.8$). Each row corresponds to a different spatial expression pattern, as illustrated in the panels (top to bottom: Pattern I-III). The diagonal dashed line with slope 1 is used as a reference.

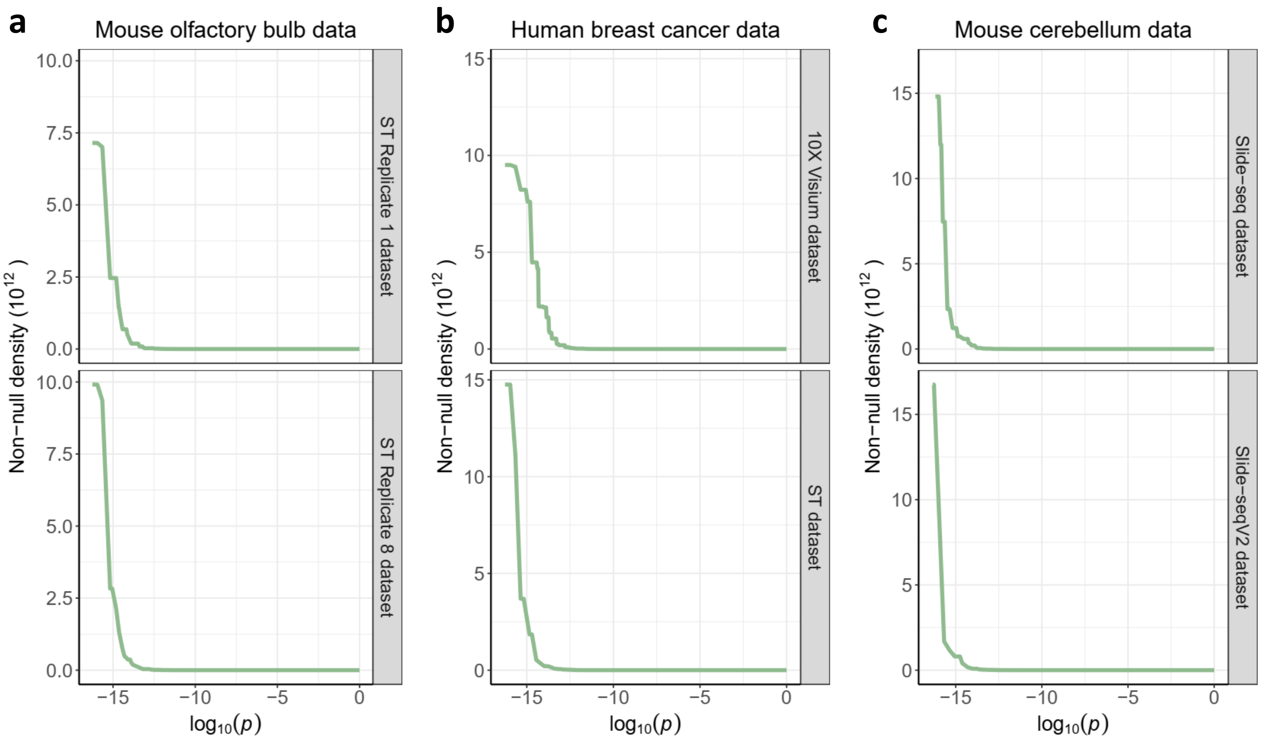


Figure S5: Plots of non-null density functions estimated by STAREG for three pairs of SRT data measured with different technologies. In each panel, the x-axis shows the $\log_{10}$ scale of $p$-values, and the y-axis shows the $10^{12}$ scale of non-null densities.

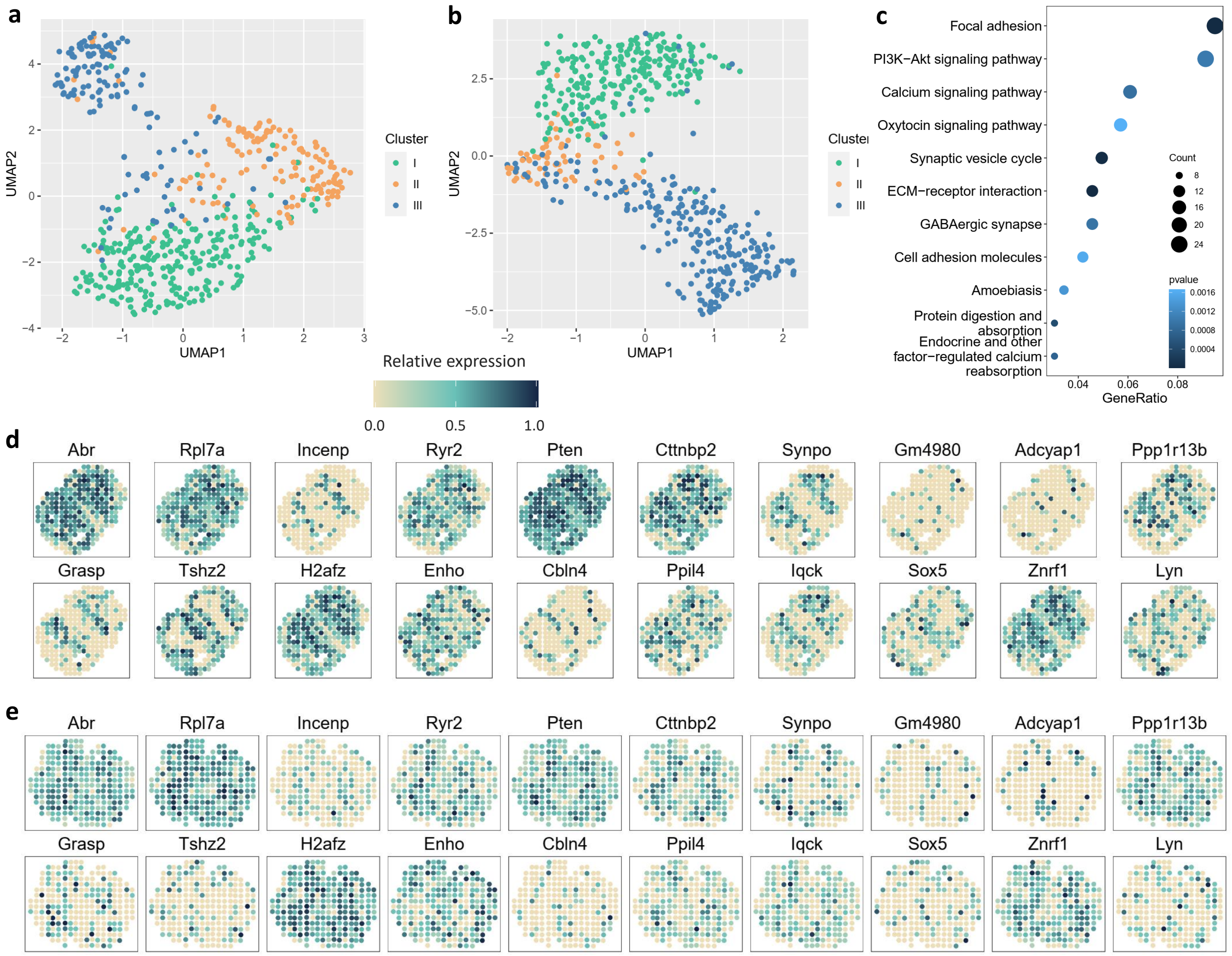


Figure S6: Analysis results of the ST data from mouse olfactory bulb.

(**a**) The Scatter plot shows spatial clusters of the ST data in Replicate 1 based on the 557 replicable SVGs only identified by STAREG. We first used UMAP to reduce the dimensionality of genes to two and then used cell labels obtained from hierarchical agglomerative clustering to visualize cell clusters.

(**b**) The Scatter plot shows spatial clusters of the ST data in Replicate 8 based on the 557 replicable SVGs only identified by STAREG.

(**c**) The bubble plot shows KEGG enrichment analysis results of the 557 replicable SVGs uniquely detected by STAREG. The color of gene sets with corresponding pathways is determined by $p$-values, and the size of bubbles represents counts.

(**d**) Spatial expression patterns of 20 genes randomly selected from the 557 replicable SVGs only identified by STAREG based on the ST Replicate 1 data. Different color represents relative gene expression levels (antique white: low; navy blue: high).

(**e**) Spatial expression patterns of 20 genes randomly selected from the 557 replicable SVGs only identified by STAREG based on the ST Replicate 8 data.

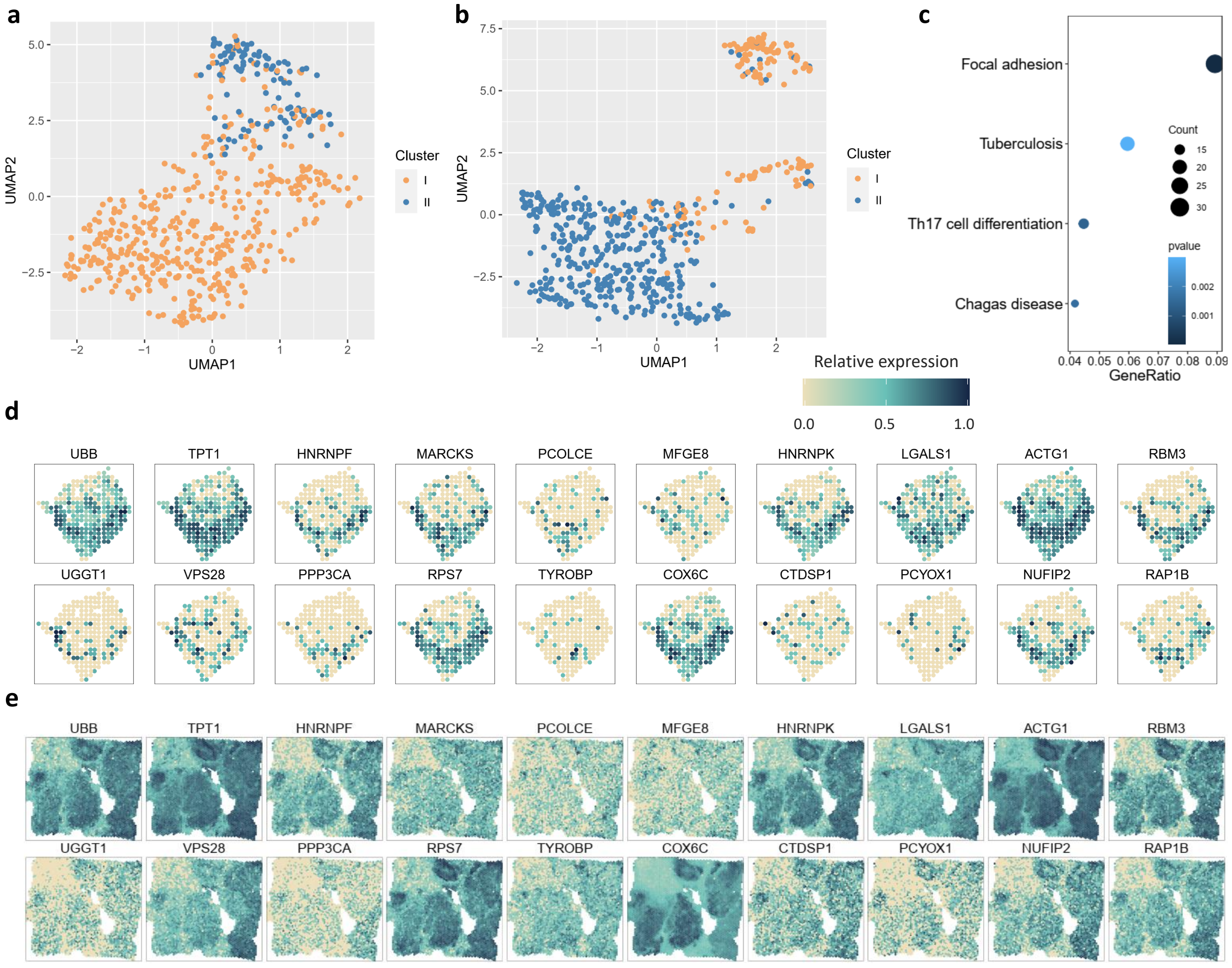


Figure S7: Analysis results of the ST data and 10X Visium data from human breast cancer.
(**a**) The Scatter plot shows spatial clusters of the ST data based on 566 replicable SVGs identified by STAREG. We first used UMAP to reduce the dimensionality of genes to two, then used cell labels obtained from hierarchical agglomerative clustering to visualize cell clusters.
(**b**) The Scatter plot shows spatial clusters of the 10X Visium data based on 566 replicable SVGs identified by STAREG.
(**c**) The bubble plot shows the 4 KEGG enrichment pathways additionally detected by STAREG. The color of gene sets with corresponding pathways is determined by $p$-values, and the size of bubbles represents counts.
(**d**) Spatial expression patterns of 20 genes randomly selected from the 62 replicable SVGs only identified by STAREG based on the ST data. Different color represents relative gene expression levels (antique white: low; navy blue: high).
(**e**) Spatial expression patterns of 20 genes randomly selected from the 62 replicable SVGs only identified by STAREG based on the 10X Visium data.

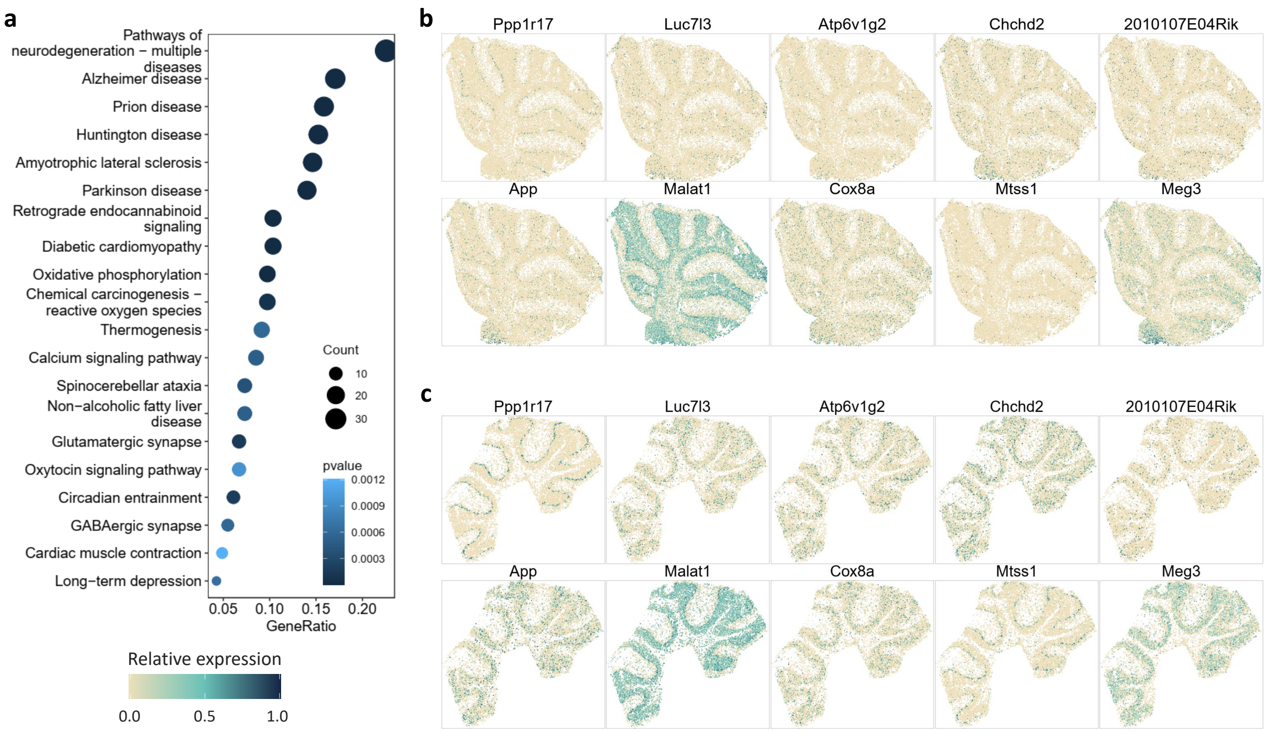


Figure S8: Analysis results of the Slide-seq data and Slide-seqV2 data from mouse cerebellum. (**a**) The bubble plot shows KEGG enrichment analysis results of the 368 replicable SVGs only detected by STAREG. The color of gene sets with corresponding pathways is determined by $p$-values, and the size of bubbles represents counts.

(**b**) Spatial expression patterns of 10 genes randomly selected from the 368 replicable SVGs only identified by STAREG based on the Slide-seq data. Different color represents relative gene expression levels (antique white: low; navy blue: high).

(**c**) Spatial expression patterns of 10 genes randomly selected from the 368 replicable SVGs only identified by STAREG based on the Slide-seqV2 data.
